# Oligomerization of peripheral membrane proteins provides tunable control of cell surface polarity

**DOI:** 10.1101/2022.01.04.474999

**Authors:** Charles F. Lang, Edwin M. Munro

## Abstract

Asymmetric distributions of peripheral membrane proteins define cell polarity across all kingdoms of life. These asymmetries are shaped by membrane binding, diffusion and transport. Theoretical studies have revealed a general requirement for non-linear positive feedback to spontaneously amplify and/or stabilize asymmetries against dispersion by diffusion and dissociation. But how specific molecular sources of non-linearity shape polarization dynamics remains poorly understood. Here we study how oligomerization of peripheral membrane proteins shapes polarization dynamics in simple feedback circuits. We show that size dependent binding avidity and mobility of membrane bound oligomers endow polarity circuits generically with several key properties. Size-dependent binding avidity confers a form of positive feedback in which the effective rate constant for subunit dissociation decreases with increasing subunit density. This combined with additional weak linear positive feedback is sufficient for spontaneous emergence of stably polarized states. Size-dependent oligomer mobility makes symmetry-breaking and stable polarity more robust with respect to variation in subunit diffusivities and cell sizes, and slows the approach to a final stable spatial distribution, allowing cells to ”remember” polarity boundaries imposed by transient external cues. Together, these findings reveal how oligomerization of peripheral membrane proteins can provide powerful and highly tunable sources of non-linear feedback in biochemical circuits that govern cell-surface polarity. Given its prevalence and widespread involvement in cell polarity, we speculate that self-oligomerization may have provided an accessible path to evolving simple polarity circuits.

**Author summary:** All cells organize their activities with respect to one or more axes of polarity. Cell polarity is often defined by the asymmetric enrichment of specific polarity proteins at the cell membrane. Absent external cues, stable polarity requires positive feedback in which proteins locally promote their own accumulation at the membrane, and the strength of feedback must depend non-linearly on local protein concentration. Here, we show that this kind of non-linear dependence arises when peripheral membrane proteins form small oligomers that dissociate from the membrane more slowly than single protein monomers. Combining this effect with a little additional linear feedback allows cells to form and stabilize asymmetric distributions of polarity proteins. In addition, we find that size-dependent reduction in oligomer mobility makes the ability to polarize more robust to variation in monomer diffusivity and cell size and makes polarity protein distributions more responsive to external inputs. Since many polarity proteins form small oligomers at the cell membrane, and there are many ways for weak linear feedback to arise in biochemical systems, the combination of oligomerization with a small amount of additional positive feedback may provide a general mechanism for polarizing a wide variety of unrelated cell types.

## Introduction

Cells rely on morphological and functional polarity to execute a wide range of biological tasks, including asymmetric cell division, polarized growth and secretion, and cell migration. Cell polarity typically emerges from underlying asymmetries in the intracellular distributions of specific molecules or molecular activities. These asymmetries can form spontaneously or in response to transient localized “symmetry-breaking” cues that determine the axis of polarity.

In many cells, asymmetries are formed at the cell surface by molecules that exchange dynamically between a rapidly diffusing cytoplasmic pool and binding sites at the plasma membrane where they undergo slower diffusion that can be further hindered by interactions with a submembrane cytoskeleton [1–3], and where they interact to promote or inhibit one another’s binding or activity [3–8]. These mutual interactions define biochemical feedback circuits, which encode the ability to respond to external cues by establishing and stabilizing asymmetric distributions of their component molecules. In the past several decades, a relatively small number of such circuits have been shown to underlie the formation and stabilization of polarity in a wide range of cellular and organismal contexts. Examples include circuits formed by small GTPases such as RhoA and Cdc42, their activators (GEFs), inhibitors (GAPs) and effectors [9, 10], the conserved PAR polarity circuit [11], the Min proteins in bacteria [12, 13], and a collection of proteins in fission yeast, including Pom1p and Mid1p, that differentiate the pole from the mid-cell [6, 7, 14].

Cell surface asymmetries can emerge spontaneously, or they can be induced by transient local cues acting in various ways – for example through locally biased activation or inhibition of recruitment [15–18] or through polarized transport [5,18,19]. However, positive feedback is required to amplify the effects of these symmetry-breaking cues and to stabilize asymmetries once the cues are gone against dissipation by dissociation and lateral diffusion. Thus, a key challenge is to understand how asymmetric distributions of membrane proteins are amplified and stabilized by the dynamic interplay of local exchange, lateral mobility, and feedback.

Theoretical efforts to address this challenge have focused on simple abstractions of known circuits or circuit “motifs” expressed as mass-conserved reaction-diffusion models [20–25]. These efforts have revealed how simple polarity circuits can manifest qualitatively different dynamics, depending upon the types and strengths of feedback, protein abundance, binding rates and mobilities, and the forms of non-linearities that appear in model equations. For example, the same model can exhibit a spatially uniform stable state, a stably polarized state, or both, depending on model parameters [25, 26]. The positions of stable domain boundaries, or the rates at which competition between multiple domains is resolved, can be continuously tuned by tuning the strength and/or saturation of feedback interactions, the lateral diffusion of proteins at the cell membrane, or the total abundance of proteins [5, 23, 27].

A general conclusion from these studies is that while linear positive feedback can generate transient noisy asymmetries when the numbers of molecules are sufficiently low [28, 29], stable polarity requires some form of non-linear positive feedback [23, 25, 30]. But how specific forms of non-linearity and feedback, arising through specific molecular interactions, shape polarization dynamics, remains poorly understood.

One potentially important source of non-linearity in polarity circuits is the oligomerization of peripheral membrane proteins. In recent years, an increasing number of polarity proteins have been observed to form discrete clusters on the cell membrane [3, 7, 31–34] suggesting that oligomerization may play a general role in polarity circuits. Oligomerization confers size-dependent membrane binding avidity [35, 36], and can also lead to size-dependent restriction of mobility through the interactions of oligomers with a submembrane cytoskeleton [3, 37–40]. A few previous modeling studies have considered oligomerization reactions [3, 6, 34, 41], but these have focused on special cases [3, 41], or on how oligomerization shapes gradients formed by a local source of protein recruitment [34, 42]. To date there has been no systematic analysis of how oligomerization of membrane proteins, and its effects on lateral mobility and dissociation, shapes their abilities to form and stabilize asymmetric distributions on the cell membrane.

Here we study a class of simple polarity models in which monomers bind reversibly to the membrane, and oligomerize in the presence of positive feedback governed by first order (linear) mass action kinetics. We find that non-linear dissociation kinetics emerge generically from size dependence of membrane binding avidity, and that this, combined with weak linear positive feedback, is sufficient for polarization. We show that the strengths of oligomerization and feedback define phase boundaries separating regimes in which stable polarity is impossible to achieve, is inducible, or occurs spontaneously. Furthermore, modulating oligomerization strength provides a simple way to tune the speed of polarization, allowing the same system to rapidly approach a uniquely stable polarized state, or to preserve polarized domains of arbitrary sizes as quasi-stable states over biologically relevant timescales. These basic findings extend to multiple circuit architectures and different modes of positive feedback. Given its widespread occurrence, our results suggest that oligomerization of peripheral membrane proteins may play a key role in facilitating and tuning polarization dynamics across a wide range of evolutionarily distinct polarity circuits and cell types.

## Results

### A kinetic model for membrane-binding and oligomerization with feedback on monomer recruitment

We consider a simple and general scenario in which monomers exchange between a well-mixed 2D cytoplasmic pool with area *A* = *HL* and a 1D membrane of length *L*, where they undergo reversible assembly into simple linear oligomers (Fig 1A). Cytoplasmic monomers also bind directly and reversibly to membrane associated monomers and oligomers. We will distinguish these two modes of binding as indirect (to the membrane) and direct (to membrane-bound monomers and oligomers). Oligomers dissociate from the membrane at a rate that decreases with oligomer size [35, 36]. To simplify the analysis, we assume that the dissociation rate is zero for oligomers with size *n* ≥2. Finally, we assume positive feedback on monomer recruitment, proportional to the total density of oligomer subunits at the membrane.

**Fig 1.**
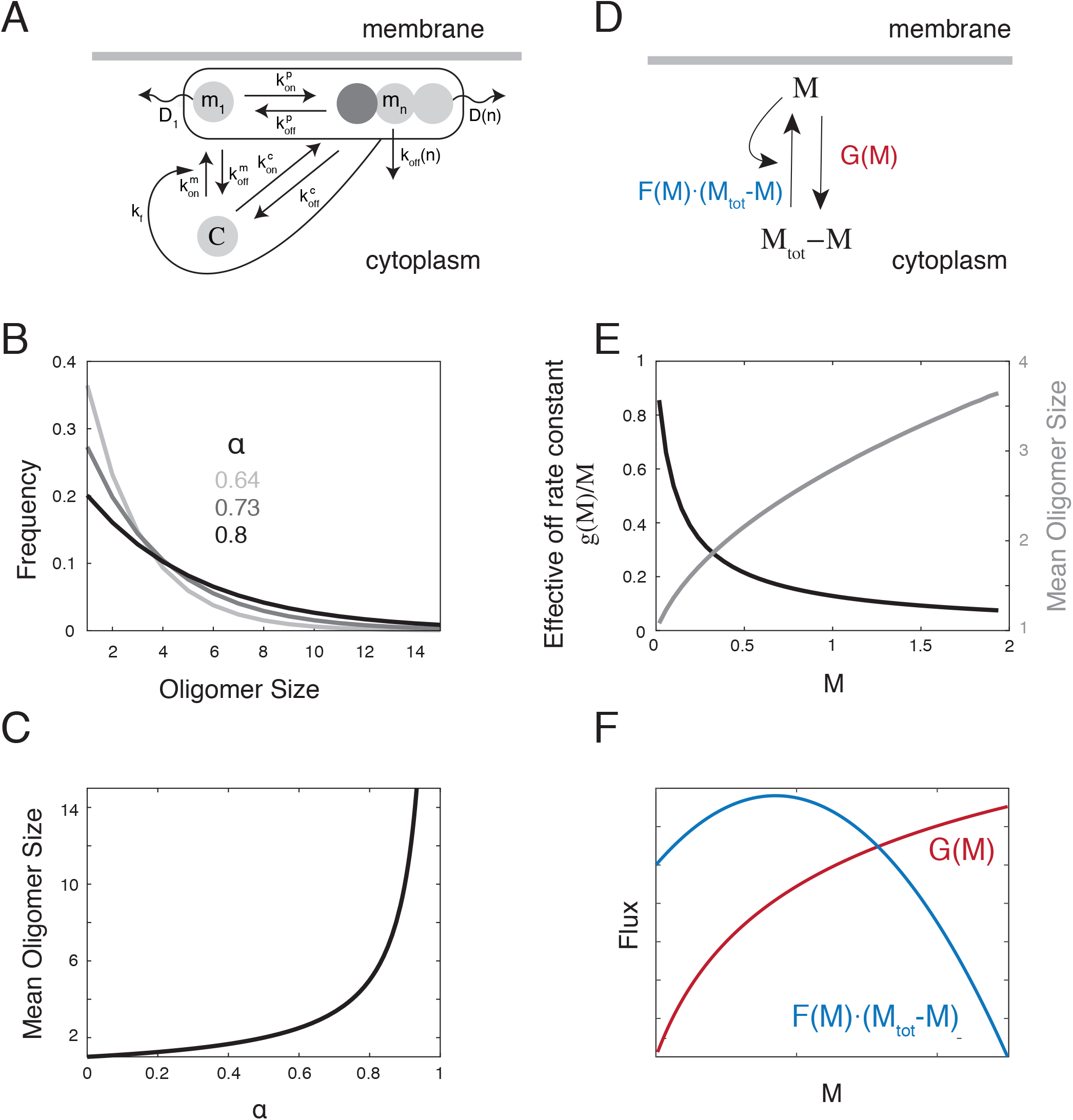
A kinetic model for membrane binding and oligomerization. **(A)** Schematic overview of the kinetic scheme. Cytoplasmic monomers bind reversibly to the plasma membrane with rate constants 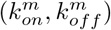, where they self-associate to form linear oligomers with rate constants 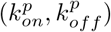. Cytoplasmic monomers also bind directly and reversibly to membrane-associated subunits with rate constants 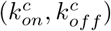. We assume that oligomer dissociation rates *k_off_*(*n*) and diffusivities *D*(*n*) are decreasing functions of oligomer size. Curved line indicates positive feedback on membrane binding at a rate *k_f_*(*M*) which depends on the local density of membrane-bound subunits *M*. **(B)** Examples of steady state oligomer size distributions corresponding to different values of the parameter *α*. **(C)** Plot showing the fixed relationship between *α* and the mean oligomer size. **(D)** A simpler model for the total density of subunits *M*, which is valid when oligomerization kinetics are sufficiently fast (see Methods for details). *F*(*M*)(*M_tot_* – *M*) and *G*(*M*) represent total binding and unbinding rates. **(E)** Effective unbinding rate (black curve) and mean oligomer size (grey curve) decrease and increase respectively as a function of *M*, reflecting size-dependent oligomer dissociation. **(F)** Representative flux balance plot showing one uniform steady state for the model. The blue line represents flux onto the membrane and the orange flux represents flux off of the membrane.

Letting x be the local position along the membrane, *m_n_* (*x*) be the density of n-mers, *N*(*x*) be the density of all oligomers, *M*(*x*) the density of all subunits, C be the concentration of cytoplasmic subunits, and *M_tot_* be the constant total number of subunits in the system, we write the following system of equations:

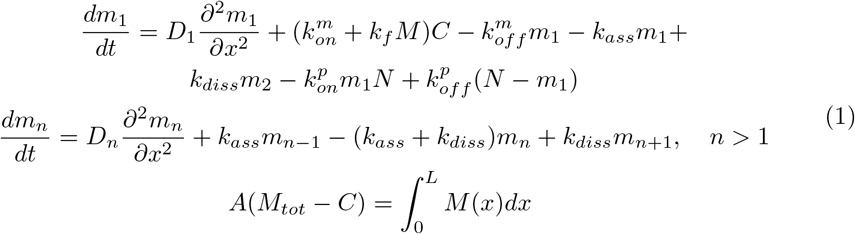

where 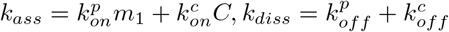 and the third equation enforces conservation of total subunits.

The spatially uniform steady states of this system are characterized by exponential distributions of oligomer sizes (see Methods) :

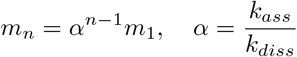

with mean cluster size:

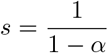

### Reduction to a one-species model

Our goal is to determine conditions in which this system will form and stabilize asymmetric distributions of membrane bound oligomers. To this end, we first consider a simpler limiting case in which oligomerization kinetics on the membrane are fast relative to the exchange of subunits between the cytoplasm and membrane. Invoking a quasi-steady state approximation to study the slower dynamics of subunit exchange, and choosing appropriate units of length, time and concentration/density (see Methods), we obtain a simpler equation for the total density of membrane-bound subunits M (see Methods):

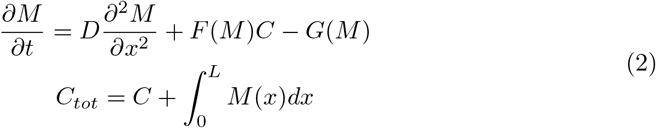

where

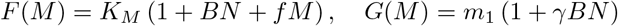

and 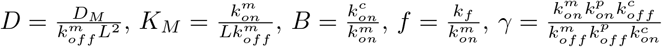 and 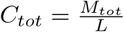 are scaled parameters. *D_M_* is the mean diffusivity of M, which will depend on mean oligomer size (see below), *B* quantifies the relative rates of direct and indirect monomer binding, *f* is the scaled feedback strength, *γ* is a dimensionless number that equals unity when the basal exchange and oligomerization kinetics (i. e. excluding feedback) (1) satisfy detailed balance, and *C_tot_* is the density of membrane-bound subunits when all subunits are on the membrane.

For this simpler system, the effective dissociation rate constant for membrane-bound subunits, 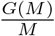, is a non-linear decreasing function of *M* (Fig. 1E, dark curve), reflecting the dependence of membrane binding avidity on oligomer size and the dependence of oligomer size on *M* (Fig. 1E, lighter curve). Thus the combination of oligomerization and size dependent binding avidity confers non-linear negative feedback on dissociation of M.

### Conditions for spontaneous and inducible polarization

Plotting binding and unbinding fluxes (*F*(*M*)(*M_tot_* – *M*) and *G*(*M*)) vs *M* shows that, for most choices of parameter values, the reduced system (and thus the full kinetic model) has a single spatially uniform steady state (Fig. 1F, S1 Fig). For different choices of model parameters, numerical solutions predict one of three qualitatively distinct behaviors (Fig 2A-C): **Spontaneous polarization** - the uniform steady steady state is unstable to all perturbations, **Inducible polarization** - the uniform steady state is stable, but stable polarity can be triggered by a sufficiently large transient perturbation, and **No polarization** - the uniform steady state is globally stable.

**Fig 2.**
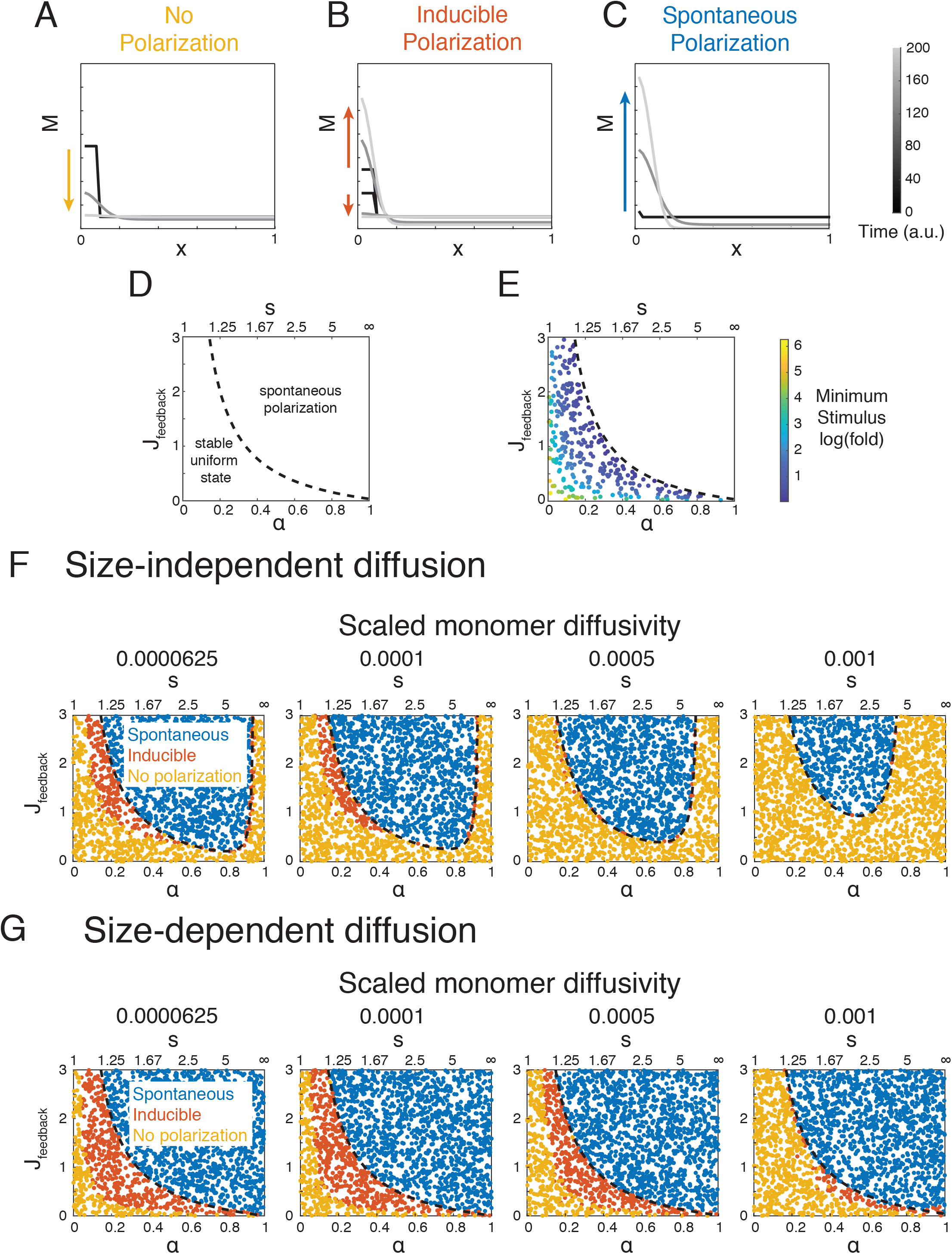
Strength of oligomerization and positive feedback define the potential for polarization. **(A-C)** Examples of three qualitatively distinct polarization regimes: **(A) No Polarization**, in which the uniform steady state is globally stable; **(B) Inducible Polarization**, in which uniform steady state is stable, but it coexists with a stably polarized state that can be accessed by a sufficiently large transient local perturbation; and **(C) Spontaneous Polarization**, in which the uniform steady state is unstable and the system will spontaneously polarize. **(D)** Spontaneous symmetry breaking as a function of *α* and mean oligomer size *s* and feedback strength *J_feedback_*. The dotted line indicates the boundary between regimes in which the uniform steady state is stable (respectively unstable) to small perturbations. **(E)** The minimum size of a local perturbation (measured as local fold increase over steady state concentration) required to induce polarization under conditions where symmetry-breaking is not spontaneous, as determined by Local Perturbation Analysis (see Methods). **(F,G)** Dependence of spontaneous and inducible polarization on scaled monomer diffusivity 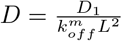. For a typical length scale *L* = 40*μm*, and monomer dissociation rate 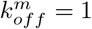, the scaled values represent (left to right) 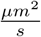. For a typical monomer diffusivity 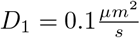, and dissociation rate 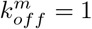, the scaled values represent (left to right) *L* = (40, 30, 20, 10) *μm*. The dotted line indicates the predicted boundary for spontaneous symmetry breaking from linear stability analysis. **(F)** represents a scenario in which diffusion is size-independent (*D_M_* = *D*_1_), while **(G)** represents a scenario in which oligomers of size greater than one do not diffuse. (*D_M_* = *D*_1_(1 – *α*)^2^).

Using linear stability analysis (see Methods), we determined general conditions for spontaneous polarization:

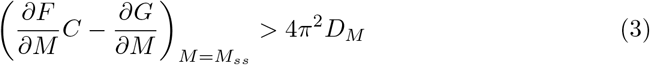

The terms on the left hand side of Eq 3 measures how the attachment and detachment rates vary with density of M near the steady state. In words, Eq 3 states that to amplify local asymmetries in M, the net accumulation rate must increase with increasing density near M, and it must do so sufficiently fast to overcome the dispersive effects of diffusion.

### Mean oligomer size and feedback strength determine the potential for polarization

We first considered the case in which diffusion is slow (*D_M_* ≈ 0) and direct binding of cytoplasmic monomers to membrane-bound oligomers can be neglected (*B* ≈ 0). Under these conditions (see Methods), spontaneous symmetry-breaking occurs when:

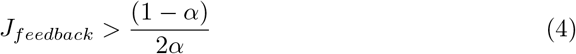

where *J_feedback_* = *fM_ss_* quantifies feedback strength as the ratio of binding flux due to feedback and the basal binding flux, and *J_feedback_* and *α* are evaluated at the uniform steady state. For 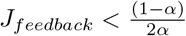, we determined the threshold for inducible polarization using Local Perturbation Analysis (LPA; [26]; see Methods). Plotting different polarization regimes in the *J_feedback_* vs *α* plane (Fig. 2D-E), highlights several conclusions: First, both indirect positive feedback (*J_feedback_* > 0) and negative dependence of dissociation rate on oligomer size (*α* > 0) are required for spontaneous symmetry-breaking. However, the strength of feedback required for spontaneous symmetry-breaking decreases rapidly with an increase in mean oligomer size, such that for mean oligomer sizes greater than a few subunits, positive feedback must only deliver a fractional increase over the basal on-rate to induce spontaneous polarization. Finally, when the uniform state is stable, stable polarity can always be induced by a sufficiently large local input, and the threshold for induction decreases with increasing *J_feedback_* or *α*, reaching zero at the spontaneous polarization boundary.

We then assessed how increasing diffusivity affects the potential for both spontaneous and inducible polarization. Recent studies suggest that the mobility of membrane-bound oligomers can decrease sharply with oligomer size [3, 34]. Therefore we considered two limiting scenarios: Size-independent diffusivity in which oligomers diffuse at the same rate as monomers (*D_M_* = *D*_1_), and size-dependent diffusivity in which oligomers of size ≥ 2 are immobile (*D_M_* = *D*_1_(1 – *α*)^2^). When diffusivity is non-negligible, the conditions for spontaneous polarization are given by

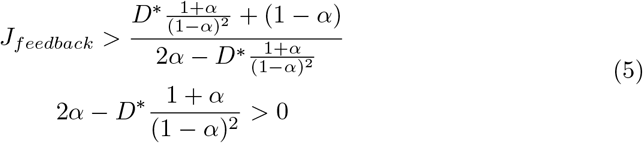

We plotted the spontaneous polarization boundary in the *J_feedback_* – *α* phase plane and used numerical simulations to assess the threshold for inducing polarity when the uniform steady state is stable (Fig. 2F,G). Consistent with intuition, we found that for size-independent diffusion, increasing scaled monomer diffusivity 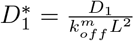 makes it harder for the system to undergo both spontaneous and induced polarization: For a given value of *α*, as monomer diffusivity increases, the feedback strength *J_feedback_* required for spontaneous polarization increases, especially for large values of *α* or oligomer size *s*, and the region of the *J_feedback_* – *α* phase plane in which polarity can be induced shrinks progressively (Fig. 2F). These effects are relatively mild for typical diffusivities of membrane proteins 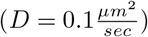, cell lengths (*L* = 10 – 40*μm*) and monomer dissociation rates 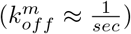 (see Fig. 2 legend for scaled diffusivities corresponding to these typical values). Importantly, for the size-dependent diffusion scenario, these effects become negligible (Fig. 2G). Thus, while diffusion can degrade the potential for spontaneous and/or induced polarization, this effect is relatively mild and it can be further reduced by a size-dependent decrease in oligomer mobility.

Numerical simulations of the full kinetic model reveal that equation 3 continues to yield an accurate prediction of spontaneous polarization when we relax the assumption that oligomerization kinetics are very fast relative to membrane exchange, i.e. when 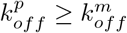 (S2 Fig). Similarly, relaxing the sharp size dependence of oligomer dissociation (S2 Fig), or allowing formation of cytoplasmic oligomers (S2 Fig), produced only minor shifts in the dependence of spontaneous and inducible polarization on *J_feedback_* and *α*. Thus simple oligomerization plus linear feedback provides a robust mechanism for spontaneous polarization of membrane bound proteins.

### Direct binding of cytoplasmic monomers to membrane-bound oligomers can promote or antagonize polarization under different conditions

In addition to binding the membrane, cytoplasmic monomers can bind directly to membrane-bound monomers and oligomers. In this case, the ability to polarize will depend on three factors: The strength of positive feedback *Jfeedback;* the ratio of direct to indirect binding rates, determined by the relative abundances of membrane binding sites and membrane bound oligomers [43], and quantified by 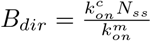, where *N_ss_* is oligomer density at uniform steady state; and whether the basal oligomerization and exchange reactions obey detailed balance.

We first asked how direct binding affects polarization driven by indirect positive feedback when basal oligomerization and exchange reactions obey detailed balance (i.e. *γ* =1) (Fig. 3A; see Methods).Neglecting the effects of diffusion, the conditions for spontaneous polarization are then given by (see Methods):

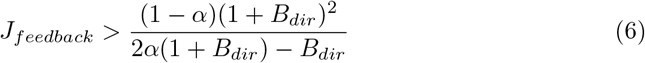

**Fig 3.**
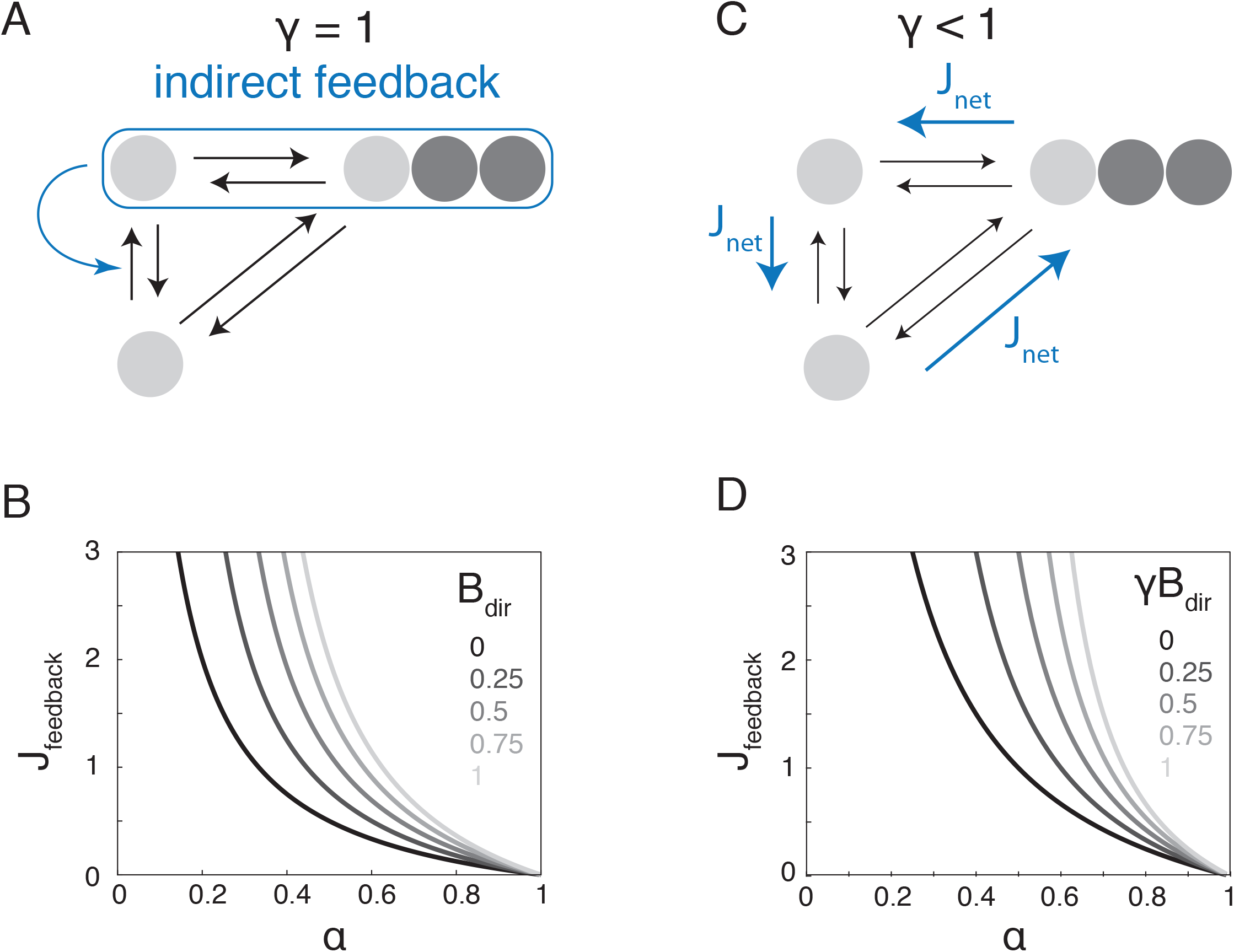
Direct binding of cytoplasmic monomers to membrane-bound oligomers can promote or antagonize polarization. **(A)** Schematic view of the scenario in which there is indirect feedback on monomer recruitment and detailed balance is enforced for membrane binding and oligomerization *γ* = 1. **(B)** Spontaneous symmetry breaking as a function of *α* or mean oligomer size *s* and *J_feedback_* for different values of *B_dir_*, measured at the uniform steady state. The grey scale lines indicate the boundary across which the uniform steady state goes from stable (bottom left) to unstable (top right) for different values of *B_dir_*. **(C)** Schematic illustrating the case where there is no indirect positive feedback and the basal kinetics obey γ < 1. **(D)** Spontaneous symmetry breaking as a function of *α* or mean oligomer size *s* and *J_feedback_* for different values of *γB_dir_*. The dotted line indicates the phase boundary separating regimes in which the uniform steady state is stable (bottom left) and unstable (top right).

Plotting the phase boundary for spontaneous polarization (Fig. 3B) shows that when detailed balance is enforced, direct binding to oligomers makes it more difficult to polarize. Increasing *B_dir_* from 0 to 1 increases the strength of positive feedback required to polarize by more than two-fold. The increase is largest when oligomerization is weak (i.e. *α* is small). Numerical simulations confirm this result for the full kinetic model when oligomerization is faster than exchange. As oligomerization kinetics become slower, the phase boundary shifts upwards in the *J_feedback_* vs *α* plane (S3 Fig). However, the effect of increasing direct binding on polarization persists. Thus direct binding to oligomers opposes polarization driven by indirect positive feedback when the basal oligomerization and exchange kinetics obey detailed balance.

We then considered an alternative scenario in which there is no indirect positive feedback (*f* = 0 in Eq 2), but one or more of the basal exchange and oligomerization reactions are driven in a way that breaks detailed balance (*γ* ≠ 1). This could arise, for example, if phosphorylation of subunits within oligomers increase their affinity for the membrane. In this scenario, spontaneous polarization can occur when *γ* < 1 (see Methods: Conditions for Spontaneous Polarization). When *γ* < 1, at steady state, there will be a net flux *J_net_* of subunits from the cytoplasm into oligomers, from oligomers onto the membrane, and then back into the cytoplasm (Fig. 3C). Defining 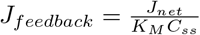 to be the ratio of this net flux to the basal rate of monomers binding to the membrane, and again neglecting diffusion, the conditions for spontaneous polarization are (see Methods):

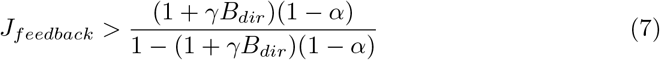

Plotting the conditions for spontaneous polarization in the *J_feedback_* vs *α* phase plane for different values of *γB_dir_* (Fig. 3D) reveals that, for relatively small mean oligomer sizes, the net flux delivered by breaking detailed balance must only be a small fraction of the basal monomer binding rate to support spontaneous symmetry-breaking (Fig. 3D). Thus when *γ* < 1, a net flux of subunits into membrane-bound oligomers constitutes a form of positive feedback. Numerical simulations show that these result hold approximately for the full kinetic model when oligomerization kinetics are sufficiently fast (S3 Fig). For slower oligomerization kinetics, and a fixed value of *γB_dir_*, there still exists a well-defined phase boundary in the *J_feedback_* vs *α* plane, but it becomes harder to polarize (S3 Fig). Altogether, these results show that when *γ* =1, direct binding opposes polarization driven by indirect positive feedback. However, when *γ* < 1, direct binding can drive polarization.

### Mean oligomer size tunes the speed of polarization and the stability of polarity boundaries

Thus far, we have characterized the conditions in which polarized states can arise through spontaneous or induced symmetry-breaking. To study how oligomerization affects the dynamic evolution of polarized states, we turned to numerical simulations. We again considered a simple model with indirect positive feedback and no direct binding, and we focused on two features of polarization dynamics: the rate at which asymmetries grow during spontaneous or induced symmetry-breaking, and the rate at which the spatial distribution of membrane-bound oligomers evolve towards a final steady state profile.

Simulations confirm that for this system, stably polarized states are characterized by single-peaked distributions (Fig. 4A). For a given choice of parameter values, the time to reach the stably polarized state depends strongly on initial conditions (S4 Fig). However, for both spontaneous and induced polarization, the growth of asymmetries proceeds through an intermediate exponential phase (Fig. 4B, S4 Fig). Although the growth rate depends on the choice of initial conditions, variation in growth rate with model parameters is tightly correlated across different initial conditions (S4 Fig). Therefore we used exponential growth rate following a transient local perturbation (3-fold increase in local protein level) of the uniform steady state as a measure of polarization speed.

**Fig 4.**
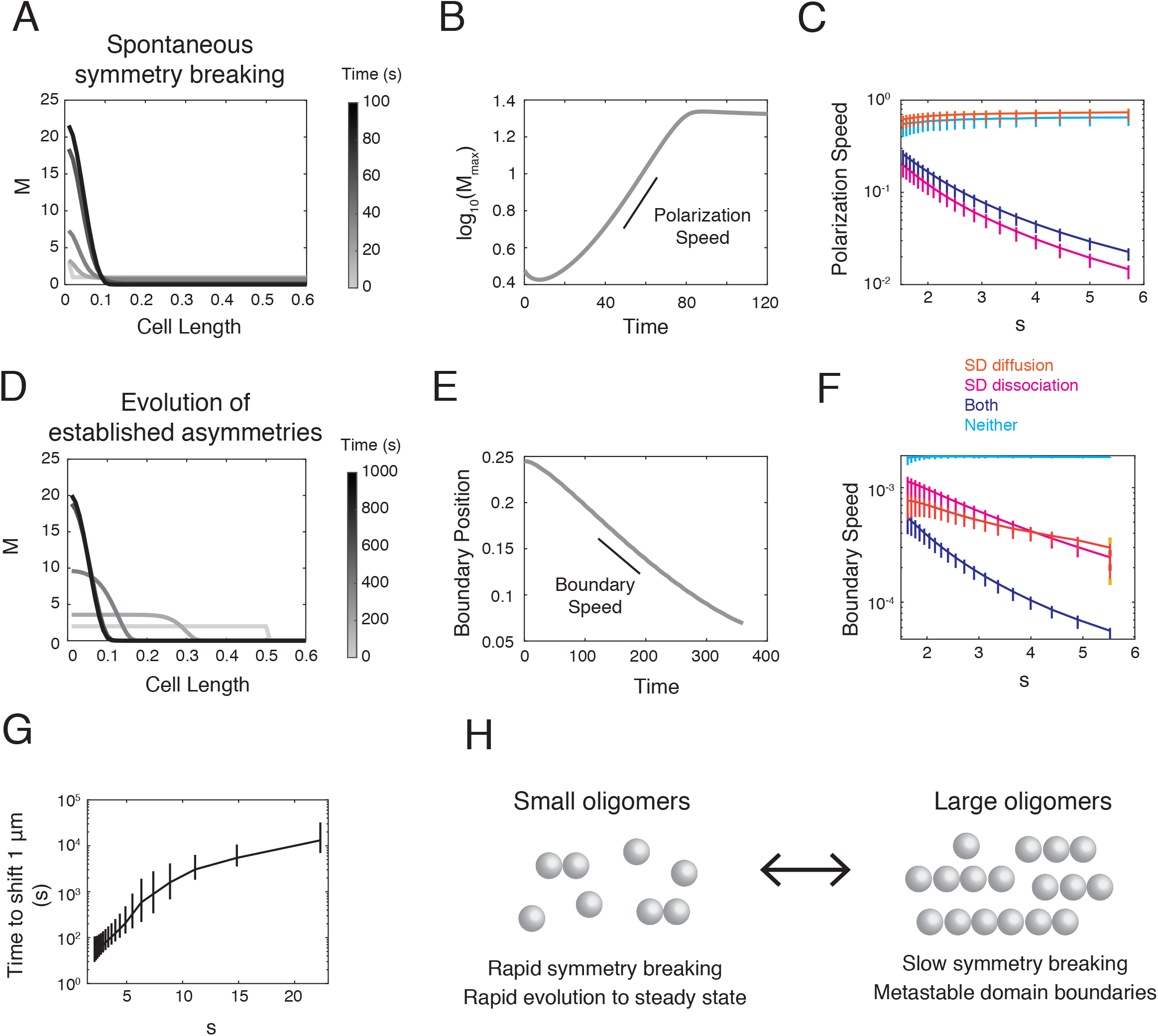
Mean oligomer size tunes polarization dynamics. **(A)** Example dynamics of polarization when symmetry-breaking is induced by a local 3-fold increase in membrane protein concentration *M* in a single point in space. Each line represents the spatial distribution of protein at a different time point **(B)** The logarithm of maximum local value of *M* over time, showing a region in which *M_max_* grows exponentially (polarization speed). **(C)** Polarization speed plotted as a function of mean oligomer size s, given different relationships between oligomer size, effective diffusivity and effective subunit dissociation rates. The bold lines indicate the average of values measured while sampling *J* between 2 and 10, and the error bars represent the maximum and minimum values. The orange line represents the case where only diffusion is size-dependent, the magenta line represents the case where only the dissociation rate is size dependent, and the dark blue line represents the case where both are size dependent. **(D)** Example of the temporal evolution of a step-change distribution that is stable in the absence of diffusion. **(E)** The boundary position, measured as the inflection point in the spatial profile of the distribution, as a function of time. The position moves at a constant rate (boundary speed). **(F)** Boundary speed plotted as a function of mean oligomer size s, given different relationships between oligomer size and effective diffusivity and effective dissociation. The bold lines indicate the average of values measured sampling *J* between 2 and 10 and the error bars represent the the maximum and minimum values. Orange line: size-dependent diffusion; Magenta line: size dependent dissociation; Dark blue line: size dependent diffusion and dissociation; Light blue line: no size dependence. **(G)** Plot showing the time it would take for the boundary to shift 1 *μm* as a function of mean oligomer size s, given *J* =1, *D* = 0.1 *μm*^2^*s*^−1^, 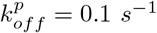, and 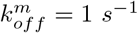. **(H)** Schematic illustrating the trade-offs that emerge from tuning the mean oligomer size.

Polarization speed was weakly sensitive to variation in feedback strength, but strongly sensitive to variation in mean oligomer size. Increasing the mean oligomer size s from 1.5 to 6 produced an order of magnitude decrease in polarization speed (Fig. 4C, blue curve; solid line represents mean speed for a given oligomer size; error bars indicate the maximum and minimum values measured for different values of J). The observed decrease in polarization speed is close to the predicted decrease in effective dissociation rate 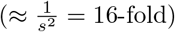, suggesting that size dependence of oligomer exchange determines how oligomerization shapes polarization speed. Indeed, scaling the monomer off-rate 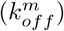 to enforce a size-independent dissociation (at steady state) completely abolished the dependence of polarization speeds on mean oligomer size (Fig. 4C, orange curve). In, contrast, nullifying size-dependence of diffusivity had no effect on polarization speed (Fig. 4C, magenta curve). Therefore, size-dependent oligomer release sets polarization speed for this simple system. Increasing mean oligomer size reduces the need for strong feedback, but at the cost of slowing down polarization.

In some biological contexts (e.g. in *C. elegans* zygotes and certain neuroblast stem cells [18, 19, 44]), a transient response to external cues can induce the rapid enrichment of polarity proteins within a broad spatial domain. Once this cue is gone, the initial distribution will evolve further through diffusion and exchange. For the system considered here, a broad initial distribution will evolve towards the stably peaked distribution described above (Fig. 4D). To determine how oligomerization and feedback shape the timescale on which this occurs, we initialized simulations with broad plateau-shaped distributions that are stable in the absence of diffusion. Then we tracked the position of the domain boundary over time in the presence of diffusion. In all such simulations, the domain boundary position moves at an approximately constant speed towards the stable peaked steady state (Fig. 4E). Therefore, we quantified how boundary speed varies with mean oligomer size (Fig. 4F). Like polarization speed, boundary speed showed very weak dependence on positive feedback and strong dependence on mean oligomer size, decreasing by more than an order of magnitude as the mean oligomer size increases from 1.5 to 6 (Fig. 4F). In this case, the size-dependence of boundary speed depends on multiplicative contributions from size-dependent dissociation and mobility (Fig. 4F). Slowing depolymerization kinetics further decreases both polarization and boundary speed (S5 Fig). For typical diffusivities of membrane proteins 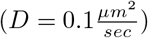, and cell sizes (*L* = 50*μm*), a mean oligomer size of 4, and 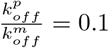 a shift in boundary position by 2 percent of cell length would take > 250 times the monomer binding lifetime, and for a mean oligomer size of 20, the same shift would take > 5000 times the monomer lifetime (Fig. 4G). Therefore, oligomerization of membrane proteins can allow a cell to effectively stabilize polarized domains of different sizes in response to transient inputs over times much longer than the residence times of individual proteins.

When boundary speed is slow relative to polarization speed, the quasi-stability of a step-change asymmetry can be assessed in a simpler two-compartment model in which two membrane compartments of varying relative lengths compete for a shared pool of cytoplasmic monomers in the absence of lateral membrane diffusion (see Methods). This analysis shows that when oligomerization and feedback strengths are tuned for spontaneous symmetry breaking (upper right domain in Fig. 2D), there is no limit to the size of the domain that a cell can stabilize. When the uniform steady state is stable (but stable polarity can still be induced; lower left domain in Fig. 2E), the maximum size of the quasi-stable domain that can be induced depends on both oligomerization and feedback strength and becomes smaller with increasing distance from the phase boundary (S6 Fig). Therefore, the ability to stabilize a domain of arbitrary size is only a property of systems in which polarization would occur spontaneously

Together, these findings reveal how size-dependent oligomer dynamics produce an intrinsic trade-off between an ability to polarize rapidly in response to small cues or noisy fluctuations, and the ability to stabilize a domain boundary at an arbitrary position (Fig. 4H). Increasing mean oligomer size pushes the system towards slow polarization and stable boundaries.

### Saturating feedback slows polarization speed but has weak effects on boundary speed relative to oligomerization

Previous studies show that introducing saturating feedback kinetics into simple mass-balanced reaction diffusion models can lead to slower polarization and slower resolution of multipolar states by competition [23]. To ask how the effects of saturation compare to those that arise through oligomerization, we modified the above model (linear indirect feedback and no direct binding) by introducing a simple form of saturating feedback proportional to 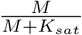 (Fig. 5A). Introducing 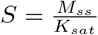 as a simple measure of saturation at the uniform steady state, the conditions for spontaneous polarization become (see Methods):

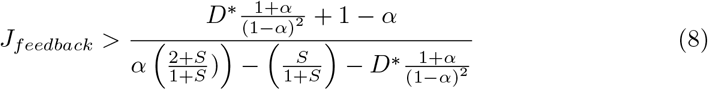

where *J_feedback_* is again the ratio of monomer binding flux due to positive feedback over the basal monomer binding rate, evaluated at the uniform steady state.

**Fig 5.**
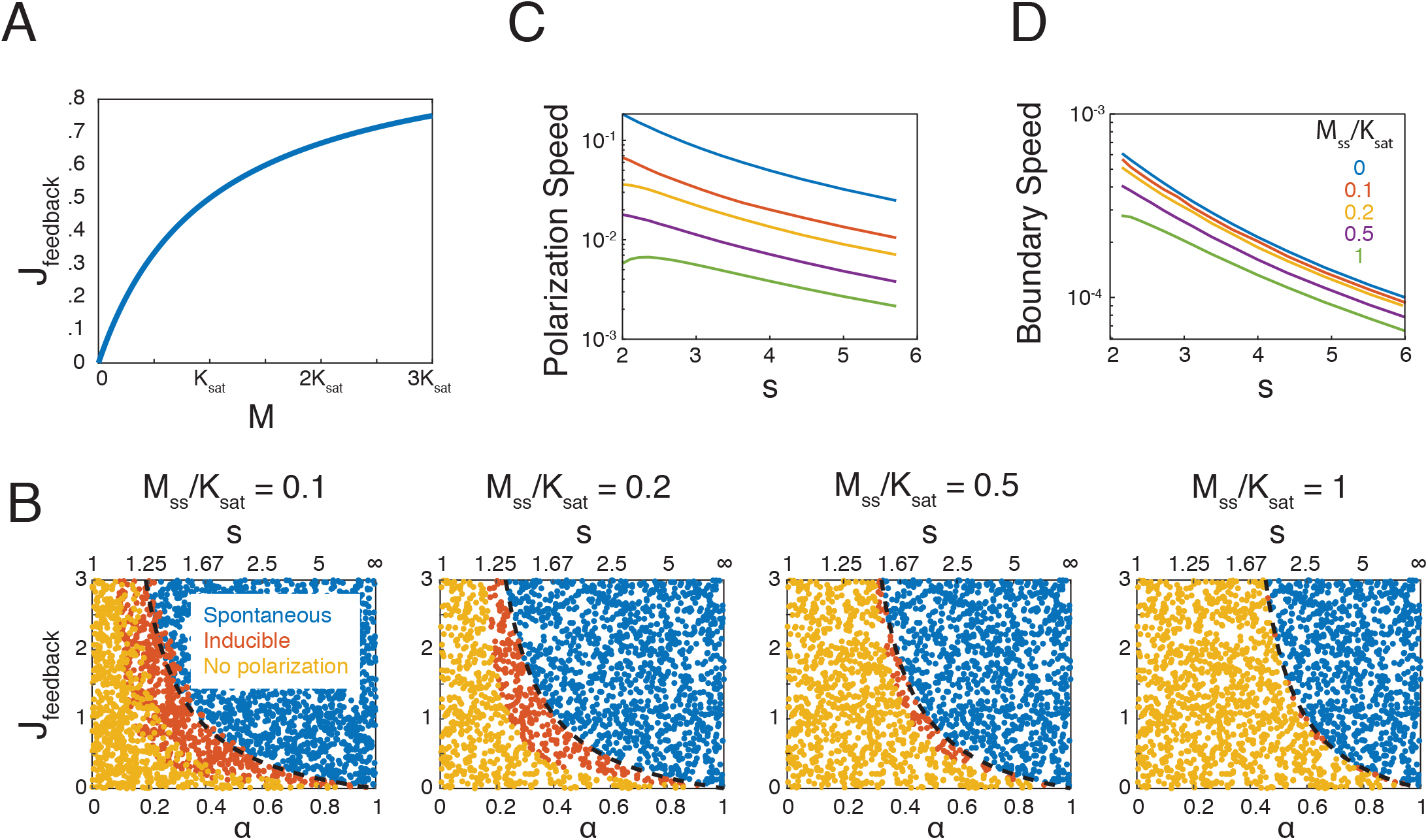
Saturating feedback reduces the potential for polarization and slows polarization speed but has weak effects on boundary speed. **(A)** Plot of simple saturating feedback. **(B)** Phase diagrams produced using LPA and showing how regimes with spontaneous, inducible, or no polarity shift with increasing saturation at uniform steady state, measured by 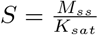. **(C,D)** The effect of increasing *M* relative to the saturation constant on the dynamics of polarization and boundary stability, assessed by numerical simulations. **(C)** Polarization speed plotted as a function of mean oligomer size s, for different values of 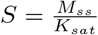. **(D)** Boundary speed plotted as a function of mean oligomer size s, for different values of 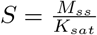.

Using a combination of local perturbation analysis and simulations, we find that introducing saturation increases the strength of feedback and/or oligomerization required for spontaneous polarization, and reduces the range of parameter values for which polarity can be induced through local perturbation (Fig. 5B). Consistent with previous reports [23], both the polarization speed and boundary speed decrease with the strength of saturation (Fig. 5C,D). However, while the effect on polarization speed is significant relative to the effects of oligomerization(Fig. 5C), the effect on boundary speed is modest(Fig. 5D). Thus increasing the degree of saturation makes it more difficult to polarize, both by increasing the strength of feedback and/or oligomerization required to polarize, and by slowing down polarization speed. In contrast, while saturating feedback makes boundaries of established asymmetries slightly more stable, its effects are relatively weak compared to those of increasing mean oligomer size.

### The role of oligomerization in shaping polarization dynamics extends to different feedback topologies

Thus far, we considered forms of feedback in which a protein acts locally to promote its own accumulation on the cell membrane. To explore the generality of these results, we considered an alternative model in which two peripheral membrane proteins *A* and *B* bind the membrane at constant rates and act locally to promote one another’s dissociation at rates that are linear functions of their local densities (Fig. 6A). We assume that *A* forms oligomers while *B* does not, and that only monomers dissociate. Assuming that oligomerization of A is fast relative to exchange, we can write equations for a two-species model:

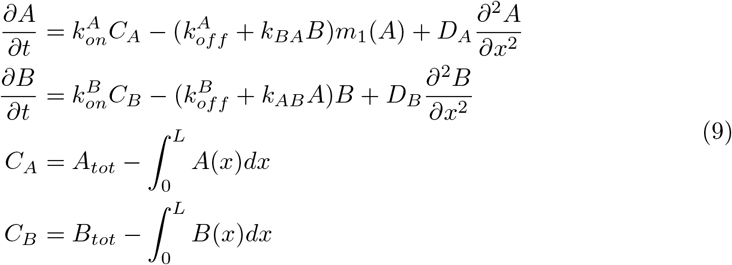

**Fig 6.**
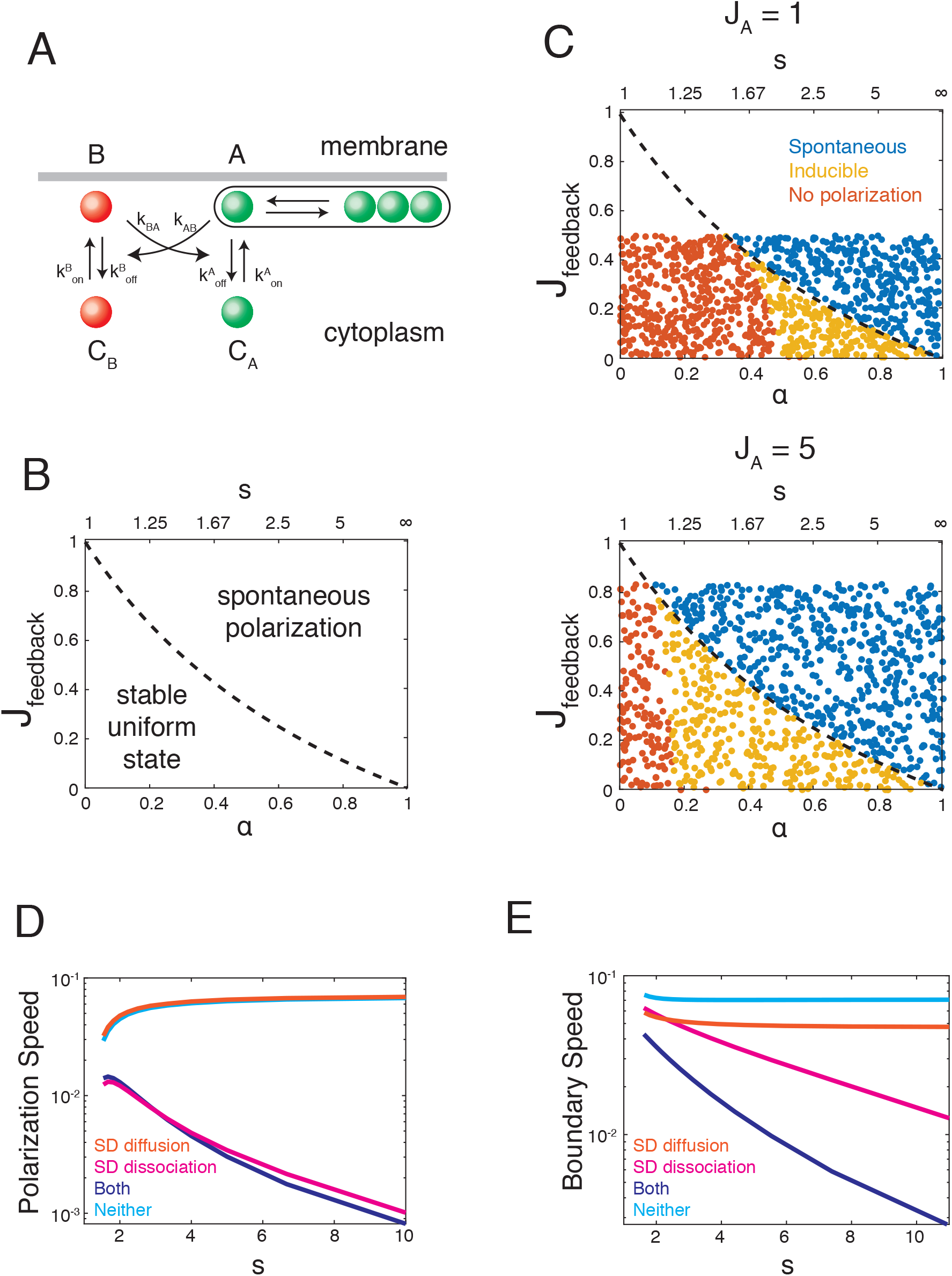
Oligomerization promotes polarization and slows polarization dynamics when combined with mutual antagonism. **(A)**Reaction diagram showing mutually antagonistic effects between two membrane-binding proteins. **(B)** Analytical solution for the boundary between unstable and stable uniform states in phase space determined by *α* and 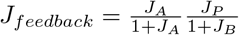. **(C)** The potential for polarization, as assessed by LPA, plotted in terms of *α* or *s* and *J_feedback_* for *J_A_* = 1 (top) and *J_A_* = 5 (bottom). Blue indicating spontaneous polarization, yellow indicating inducible polarization, and orange indicating no polarization. **(D)** Polarization speed plotted as a function of mean oligomer size *S*, given different relationships between oligomer size and effective diffusivity and effective dissociation. The orange line represents the case where only diffusion is size-dependent, the magenta line represents the case where only the dissociation rate is size dependent, and the dark blue line represents the case where both are size dependent. **(E)** Boundary speed plotted as a function of mean oligomer size *S* for *J_feedback_* = 0.64, given different relationships between oligomer size and effective diffusivity and effective dissociation. Orange line: size-dependent diffusion only; Magenta line: size dependent dissociation only; Dark blue line: size dependent diffusion and dissociation; Light blue line: no size dependence.

Considering a limiting form of this model in which diffusion is slow and *B* monomers dissociate far more rapidly than *A* monomers, we find that spontaneous polarization occurs when:

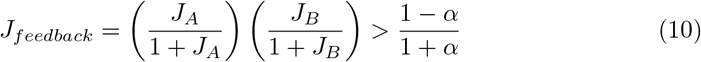

where *J_A_* and *J_B_* are the ratios of feedback-dependent to basal dissociation rates for *A* and *B* respectively. Plotting *J_feedback_* vs *α* (Fig. 6B) shows that as for models with positive feedback on local recruitment, combinations of relatively weak oligomerization and feedback are sufficient for spontaneous polarization. For example, when the mean oligomer size is 2.5, mutual inhibition must only double the dissociation rates of *A* and *B* for polarization to occur. Local perturbation analysis reveals that even when polarization does not occur spontaneously, it can be induced by a sufficiently large local perturbations for a broader range of *J_feedback_* and *α* values (Fig. 6C). Finally, as for models with positive feedback on recruitment, the polarization speed and the boundary shift speed decrease as a function of mean oligomer size (Fig. 6D). Thus, oligomerization enables polarization and shapes polarization dynamics in very similar ways for networks with very different feedback topologies.

## Discussion

The ability to self-oligomerize is common to many peripheral membrane proteins that adopt asymmetric distributions in polarized cells. Here we have explored how this ability shapes the performance of simple feedback circuits that underlie cell polarization. We focused on two properties that emerge as natural consequences of oligomerization – size-dependent membrane binding avidity [35, 36] and size-dependent mobility [40]. We find that both properties sharply enhance the ability of simple circuits to form and stabilize cell surface polarity. Both also contribute to controlling the rate at which asymmetries grow spontaneously or in response to local cues and the rate at which they evolve towards stably polarized states. Overall, our results reveal how strength of oligomerization, which determines mean oligomer size, can act as a physiological control parameter that determines if, when and how fast polarization can occur.

### Oligomerization promotes symmetry breaking through positive feedback and by reducing the dissipative effect of diffusion

Cell surface asymmetries are formed and maintained by the local differences in the relative rates of binding and unbinding [24] working against the dissipative effects of diffusion. Theoretical studies have established that both spontaneous emergence and maintenance of stable asymmetries require non-linear positive feedback on accumulation [23, 25, 30]. For peripheral membrane proteins, this can involve either increased binding rates, or decreased unbinding rates, with increasing density. To amplify local asymmetries around a spatially uniform steady state, the net rate of accumulation must increase with cell surface density at the steady state (see Eq. 3, [23, 25]). For a typical protein that binds the membrane with zero-order (density-independent) kinetics and unbinds with first order (linear in density) kinetics, this implies that the combination of positive feedback on binding and detachment must contribute stronger than linear dependence of net accumulation rate on density, to amplify local asymmetries.

Here, we find that size dependent membrane binding avidity endows oligomers generically with a form of positive feedback in which the effective dissociation rate constant decreases with increasing density. This form of feedback is insufficient, on its own, to drive spontaneous symmetry-breaking. However, it strongly reduces the amount of additional positive feedback required to break symmetry to an extent that increases with increasing oligomer size. Even for weak oligomerization, characterized by a mean oligomer size of 2.5, positive feedback on monomer binding (resp. unbinding) need only deliver an approximately 50% increase (resp. decrease) over basal rates to drive symmetry breaking. As a simple comparison, achieving the same result (with the same feedback strength) with cooperative positive feedback on monomer recruitment would require greater than fourth order dependence S7 Fig.

We also find that the size-dependent decrease in oligomer mobility can sharply reduce the dissipative effects of diffusion which would otherwise degrade the potential for symmetry-breaking. This effect would be weak for the weak size-dependence of proteins observed in pure membranes. However, interactions with a submembrane cytoskeleton can lead to much sharper size dependence, and thus much sharper reduction in mobility. Moreover, the magnitude of this effect will increase with the strength of oligomerization. Thus oligomerization of peripheral membrane proteins provides a tunable form of feedback, and tunable control of protein mobility that can greatly enhance the potential for symmetry breaking and polarization. Importantly, the strength of this affect can be readily controlled by modulating the equilibrium binding constant for self-association, e.g through phosphorylation.

### Direct binding of cytoplasmic monomers to oligomers can either inhibit or drive polarization

For proteins that can bind membranes and self-oligomerize, there are two paths by which a cytoplasmic monomer can join membrane-bound oligomers – either indirectly by first binding the membrane and then diffusing into contact with an oligomer, or by direct binding to the oligomer. Classical studies [43, 45] have characterized the relative contributions of these two binding modes to protein absorption on membranes [45] or to ligand binding and uptake [43].

Here we characterized the relative contributions of direct and indirect biding modes to polarization. We find that the contribution of direct binding depends on whether or not the basal membrane binding and oligomerization kinetics obey detailed balance (i.e. whether or not they consume energy). When detailed balance is satisfied and polarization is driven by positive feedback on monomer binding to the membrane, direct binding to oligomers reduces the potential for polarization. However, another unanticipated mode of feedback emerges when detailed balance is broken in a way that favors a steady state flux of monomers from the cytoplasm into oligomers. This could occur in many ways. For example, oligomers could recruit an enzyme that locally modifies monomers or phospholipids to promote association between monomers and the cell membrane. Regardless of how it arises, net flux from cytoplasm into oligomers at steady state is a form of positive feedback because its magnitude increases with overall density of membrane-bound subunits. As with other forms of feedback, we find that the potential for polarization increases with feedback strength (measured as the ratio of the net flux from cytoplasm into oligomers (*J_net_*) over basal monomer binding rate), and increasing oligomerization strength reduces the amount of additional feedback required for polarization. However, in this case, increasing direct binding increases feedback strength because it increases (*J_net_*).

Our results therefore highlight two general modes of positive feedback on recruitment that could drive symmetry breaking and polarization: One relies on increasing the effective number of membrane binding sites for cytoplasmic monomers; the other relies on breaking detailed balance to favor a net flux into oligomers. Because binding to the cell surface is a diffusion limited process, the relative densities of membrane binding sites and oligomers will determine which of these two modes are more effective drivers of cell polarity.

### Oligomerization provides tunable control over the speed and mode of polarization

Our results also reveal how oligomerization allows tunable control over the speed of polarization. Weak oligomerization favors rapid growth of asymmetries and a rapid approach to a steady state in which the position of polarity boundaries are dictated by binding/unbinding kinetics and diffusivities intrinsic to the polarity circuit through mechanisms such as wave-pinning [22]. Stronger oligomerization reduces the rate at which local asymmetries grow, and the rate at which they approach steady state. This allows external cues that promote local binding [14, 17, 46, 47] or unbinding [48] of polarity factors, or their rapid transport by actomyosin flows [18, 19, 44] to impose spatial asymmetry patterns that are then maintained as quasi-stable states over much longer timescales, and which are no longer dictated by the internal reaction/diffusion kinetics of the polarity circuit. While increasing oligomerization strength to maintain asymmetries far from steady state will also necessarily decrease the rate at which asymmetries grow, this can be readily overcome through the rapid control of local binding/unbinding or transport by external inputs. Thus, modulating the strength of oligomerization can allow tunable control, across different cells or within the same cell over time, over the relative extents to which circuit-specific reaction diffusion dynamics and external inputs determine the spatial distributions of polarity proteins.

### Homo-oligomerization as a versatile source of non-linear feedback and tunable mobility for cell polarity

The ability to self-oligomerize is common to a large fraction of peripheral membrane proteins. By recent estimates, something like 50% of cytoplasmic and membrane proteins form homo-oligomers [49, 50], although the majority of these form homo-oligomers of fixed size. This abundance is thought to be driven at least in part by the ease with which homodimeric interfaces can arise through random mutation and through selection for the many functional advantages (unrelated to polarization) conferred by homo-oligomerization [49, 51], and also through entrenchment of randomly occurring, selective neutral mutations [52]. Therefore, the frequent presence of oligomeric proteins in polarity circuits may be the result from exaptation of a frequently occurring property of proteins for an essential function in polarization. Importantly, because the generic form of non-linear positive feedback and tunable control of mobility conferred by oligomerization do not depend on polarity circuit architecture, or other modes of feedback, the barrier to exaptation is low.

## Methods

### Mathematical Model

We consider a simple 2D “cell” consisting of a 1D membrane of length L adjacent to a cytoplasmic compartment with height H and area A = HL. Without loss of generality, we assume *H* = *L*. We assume that monomers bind reversibly to the membrane from a single well-mixed cytoplasmic pool with rate constants 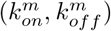, and they self-associate at the membrane with rate constants 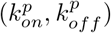 to form simple linear oligomers. In addition, cytoplasmic monomers can bind directly and reversibly with rate constants 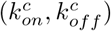 to monomers or the ends of membrane-bound oligomers. We assume that there is simple positive feedback on monomer recruitment, proportional to the total density of oligomer subunits at the membrane. Finally, we assume that oligomers dissociate from the membrane at a rate that decreases with oligomer size. We assume initially that oligomers with size > 2 are stably bound.

Letting:

*m_n_*(*x*) be the concentration of oligomers of size i,
*N*(*x*) be the total density of oligomers at the membrane,
*M*(*x*) be the total density of oligomer subunits at the membrane
*C* be the total concentration of cytoplasmic subunits,
*M_tot_* be the total number of all oligomer subunits

we write the following system of equations

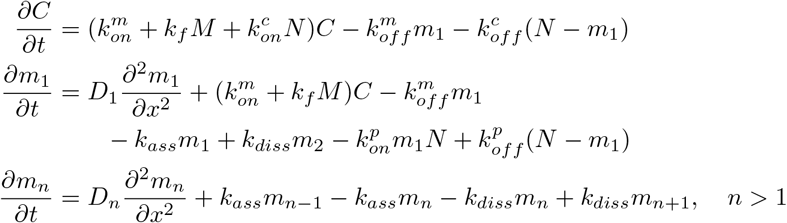

where 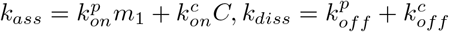. Note that these equations are not independent, given the conservation of total subunits:

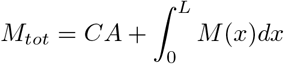

### Steady state analysis

Setting time and space derivatives to 0 in Eq 11, we obtain:

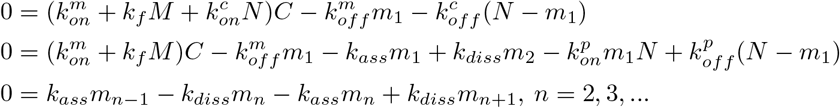

The bottom equations imply:

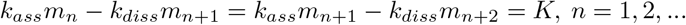

Combining the top two equations yields:

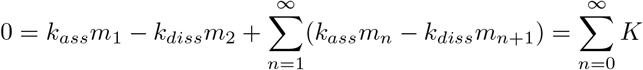

It follows that *K* must be identically 0, and at any uniform steady state, we must have

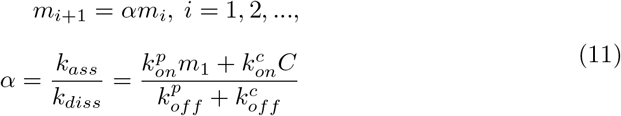

Summing 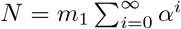 and 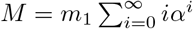 yields the following simple identities relating *m*_1_, *N*, *M* and *α* that we use below:

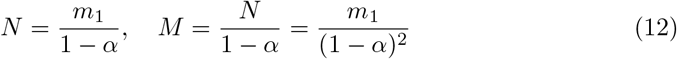

### Enforcing detailed balance for basal binding kinetics

If basal membrane binding and oligomerization reactions satisfy detailed balance, then the free energy change associated with incorporating a cytoplasmic monomer into a membrane-bound oligomer is path-independent:

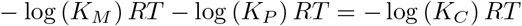

where

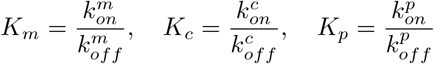

and therefore detailed balance is satisfied when:

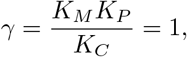

### Reducing to a simple one species model

We start by rewriting:

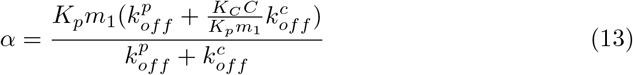

If oligomerization kinetics are sufficiently fast, i.e if they satisfy:

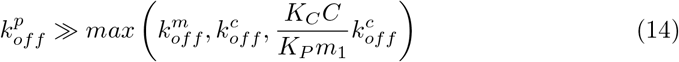

Then *α* ≈ *K_p_m*_1_, and we can invoke a quasi-steady state assumption to write the single equation for *M*:

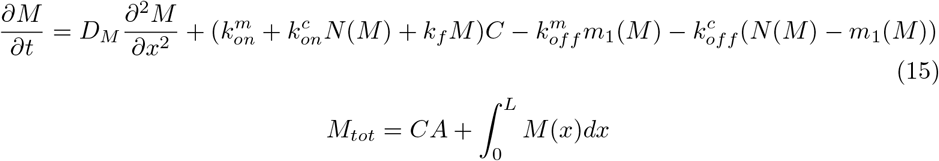

where

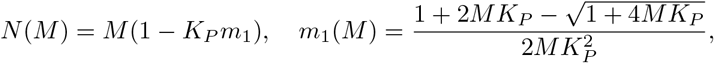

and where *D_M_* represents the average subunit diffusivity, reflecting the dependence of oligomer mobility on oligomer size.

Rearranging terms and using *N* – *m*_1_ = *K_P_m*_1_*N*:

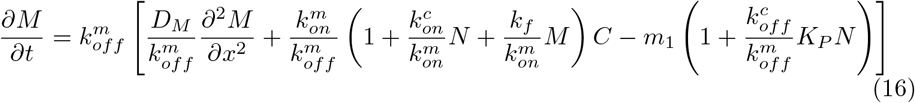

Choosing units of time 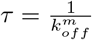, length *l* = *L* and membrane density *ρ* we obtain:

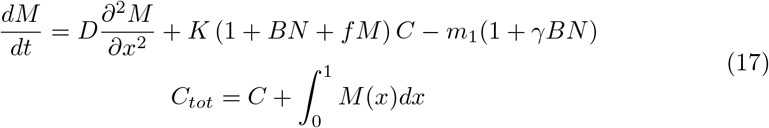

where:

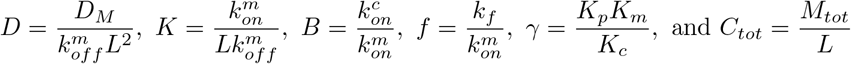

### General conditions for spontaneous polarization

Eq 17 has the general form:

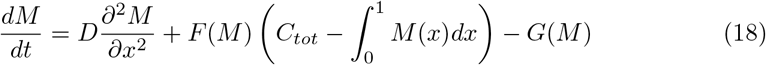

where *F*(*M*) and *G*(*M*) define membrane binding and unbinding kinetics, respectively, as a function of *M*. Spatially uniform steady state solutions of the form *M*(*x*) = *M_ss_* must satisfy:

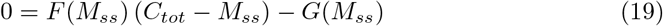

We consider stability with respect to small perturbations of the form:

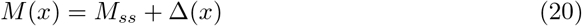

Any such perturbation can be written as 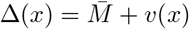, where 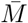 is a spatially uniform perturbation and *v*(*x*) is a spatially varying perturbation satisfying

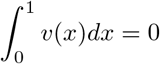

Combining Eqs 18 and 19:

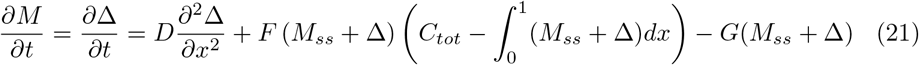

Using a Taylor’s series expansion, we obtain:

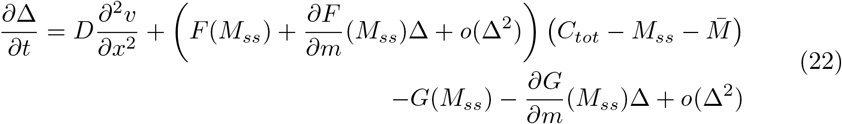

Neglecting higher order terms and using

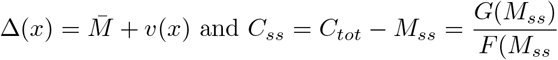

we have:

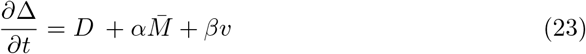

where:

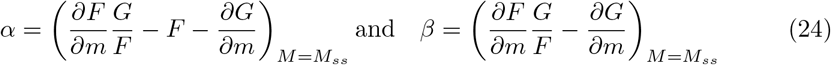

The general solution to Eq 23 can be written:

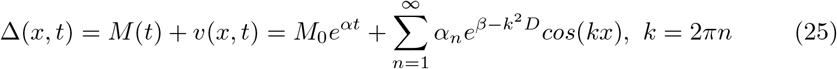

For any small perturbation 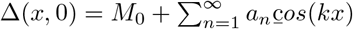, the spatially uniform part, *M*(*t*) = *M*_0_*e^αt^*, decays over time if *α* < 0, whereas the spatial differences 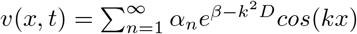, can grow only if *β* > *k*^2^*D* for some *n* ≥ 1. Therefore, the conditions for spontaneous polarization are:

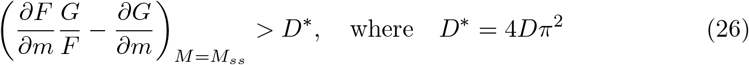

### Conditions for inducible polarization

We used Local Perturbation Analysis (LPA) to evaluate coexistence of stable spatially uniform and polarized steady states. Briefly, we consider the dynamic response to a finite-amplitude perturbation from a stable spatially uniform steady state *M* = *M_ss_* within an infinitesimally small spatial domain. We assume that diffusion is sufficiently slow (*D_M_* ≈ 0) that local changes within this domain do not affect the bulk pools of membrane-bound or cytoplasmic protein. The equation for the local response *M_l_* then writes:

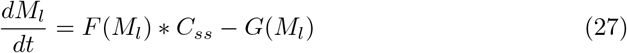

where

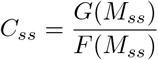

Eq 27 has a fixed point at *M_l_* = *M_ss_* corresponding to the spatially uniform steady state. When *D_M_* = 0, spontaneous polarization occurs when this fixed point is unstable, i.e when:

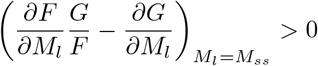

consistent with the result from linear stability analysis when *D_M_* = 0.

If the fixed point at *M_l_* = *M_ss_* is stable, then a stably polarized steady state exists when Eq 27 has an unstable steady state at *M_l_* = *M_th_*, for *M_th_* > *M_ss_*. *M_th_* then defines the threshold size of a perturbation required to induce a transition to the polarized steady state (neglecting diffusion).

In practice, we used MATLAB to identify a point *M_l_* = *M_th_* > *M_ss_* where:

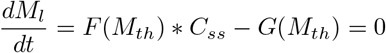

within the range:

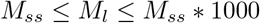

and we assessed its stability by determining the sign of:

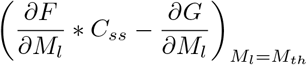

### Specific conditions for spontaneous polarization

#### Case 1: Indirect positive feedback; basal kinetics satisfy detailed balance

If basal kinetics satisfy detailed balance, then *γ* =1,

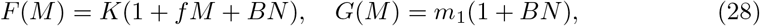

and the uniform steady state is unstable when

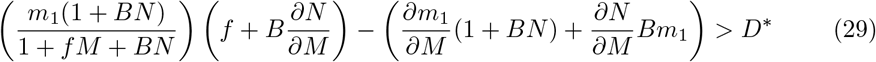

where *m*_1_, *N* and *M* are evaluated at the uniform steady state.

Using:

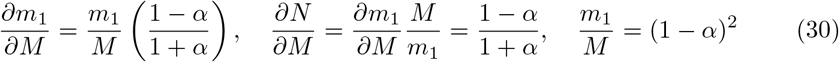

gives:

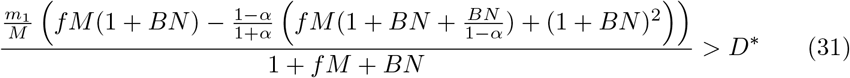

Introducing *J_feedback_* = *fM*, and *B_dir_* = *BN*, this becomes:

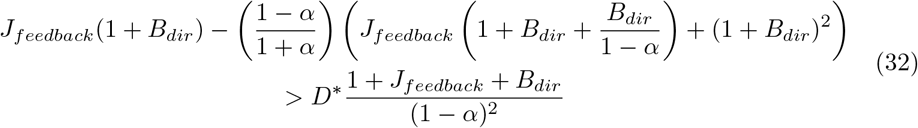

Solving for *J_feedback_* yields conditions for spontaneous polarization:

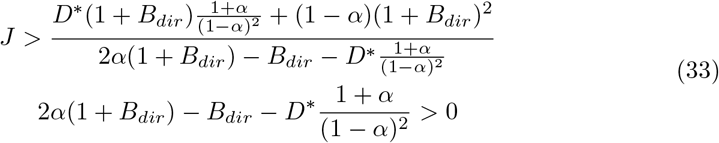

The simpler conditions cited in the main text follow directly from setting *D*\* =0 and/or *B_dir_* = 0 in Eq 33

#### Case 2: No indirect positive feedback; detailed balance not satisfied

Assume now that there is no positive feedback (*f* = 0) and that detailed balance is not satisfied (*γ* ≠ 1):

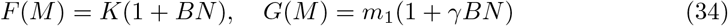

The uniform steady state is unstable when

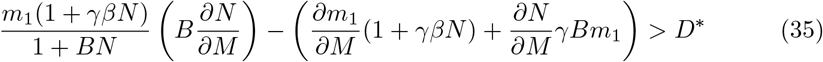

Using

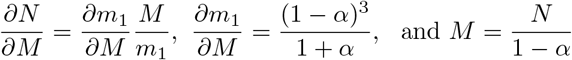

and simplifying yields:

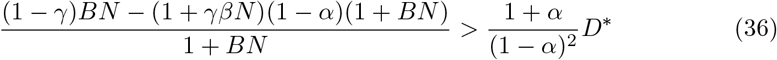

Note that this inequality can only be satisfied if *γ* < 1. Thus detailed balance must be broken in a particular direction for spontaneous symmetry breaking to occur.

To simplify this inequality, we consider the net flux of monomers from the cytoplasm to membrane oligomers (or from the membrane back to the cytoplasm) at steady state. From eq 17, we have:

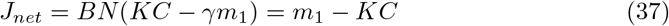

Eliminating *KC* and solving for *J_net_* yields:

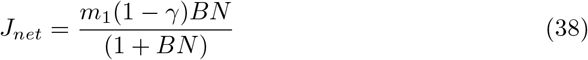

or

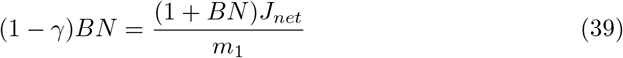

Using *J_net_* = *m*_1_ – *KC* and introducing 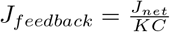, this becomes:

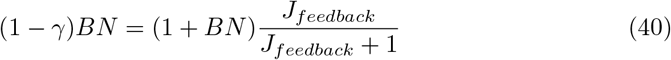

Substituting 40 into 36, and using *B_dir_* = *BN* gives:

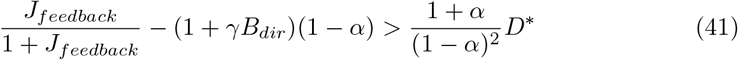

Solving for *J_ratio_* yields the final result:

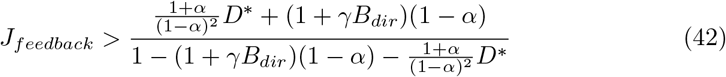

When *D*\* =0, this reduces to the simpler form:

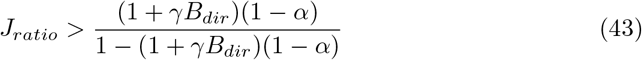

#### Case 3: Saturating feedback and no direct binding

We now consider a model in which there is no direct binding and positive indirect feedback on M recruitment is governed by a saturating function of the form:

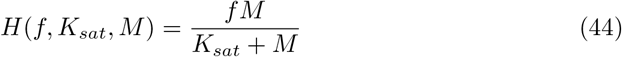

With these assumptions, we have:

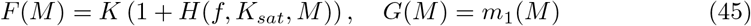

Thus the conditions for spontaneous polarization are:

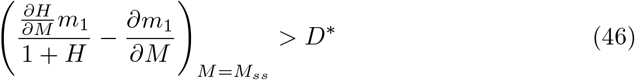

One can readily verify that::

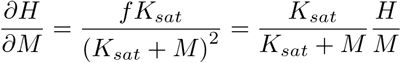

Using

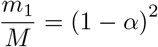

and introducing:

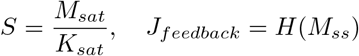

we have:

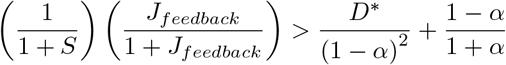

Finally, solving the inequality for *J_feedback_*, we find that spontaneous polarization occurs when:

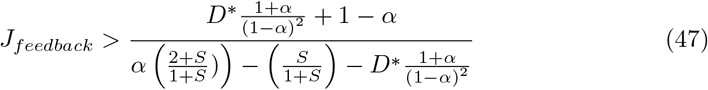

#### Case 4: Feedback through cross-inhibition

We consider a model in which two distinct proteins *A* and *B* bind reversibly to the membrane from well-mixed cytoplasmic pools. We assume that *A*, but not *B*, oligomerizes at the membrane, and that cytoplasmic monomers of *A* do not bind directly to membrane-bound oligomers. We assume that *A* and *B* monomers bind to the membrane at constant rates, and that *A* promotes the dissociation of the *B* monomers, and *B* promotes the dissociation of *A* monomers, in each case at rates proportional to the total density of membrane bound protein. Assuming that oligomerization of *A* is much faster than exchange, and invoking a quasi steady state approximation described above, we can write:

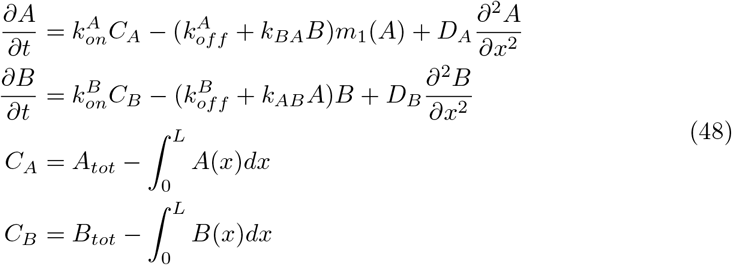

Where *k_AB_* and *k_BA_* are cross-inhibition rates and:

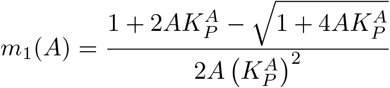

One can verify that this system of equations has a single spatially uniform steady state. Assuming that diffusion of *A* and *B* is negligible, we can determine the conditions for spontaneous symmetry breaking by considering the dynamic response of *A* and *B* to a local perturbation to this steady state within an infinitesimally small domain. The time evolution of local densities of *A* and *B* are then given by:

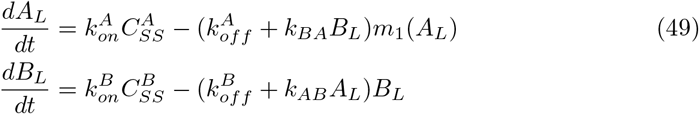

Assuming that the membrane-binding dynamics of *B* are much faster than those of *A* and making a quasi-steady state approximation for *B_L_*:

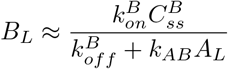

we obtain:

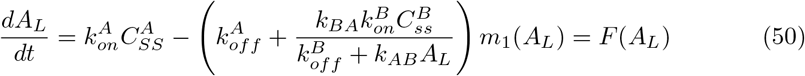

Eq 50 has a steady state at *A_L_* = *A_ss_*. If diffusion is infinitely slow and membrane dynamics of *B* are infinitely fast, a small perturbation to the steady state in equations 48 will grow iff this steady state is unstable:

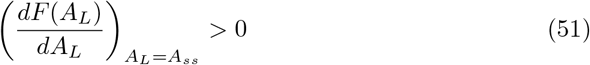

Thus we have:

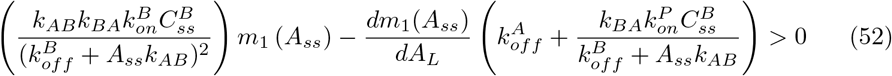

using

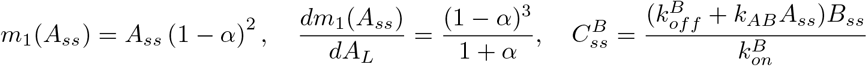

we have:

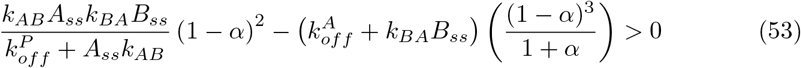

and thus:

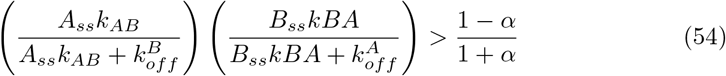

Finally, introducing 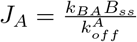 and 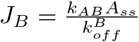 we obtain the final result:

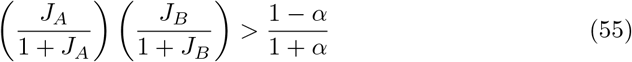

### Numerical simulations

We performed all numerical simulations using custom scripts written in MATLAB. We discretized the 1D membrane domain into 100 equally-sized compartments and used a center-difference approximation to estimate diffusive flux between compartments. We used periodic boundary conditions in simulations to test for symmetry breaking, and no-flux boundary conditions in simulations to determine dynamics. In simulations of the full kinetic model, We allowed oligomers a maximum size of 50. We solved the resulting systems of ordinary differential equations using MATLAB’s built-in ode45 function using default settings. In all simulations, we chose 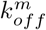 and membrane length *L* to be the units of time and length respectively, and selected values for other model parameters as described below.

### Mapping the parameter space for spontaneous and inducible polarization in simple one- and two-species models

#### Linear positive feedback and no direct binding

For this scenario, we set *B* = 0 and *γ* = 1 in Eq 17. We randomly sampled values for *K_M_* between 0.2 and 2, *α* between 0 and 1 and *J_feedback_* between 0 and 3. We constrained the uniform steady state density to be *M_ss_* = 1, and then we computed the values of all other model parameters. For the case with no diffusion (*D* = 0, Fig. 2E), we used LPA as described above to determine the threshold for inducible polarization. For the case where *D* > 0 (Fig. 2F), we chose values for *D* as specified in the figure legend, sampled all other parameter values as described above, and then assessed the potential for spontaneous and inducible polarization using simulations. To look for spontaneous polarization, we initialized simulations from the uniform steady state with a small perturbation to the density in one membrane compartment. To look for polarity induction, we initialized simulations with an asymmetry predicted to be stable without diffusion based on phase plane analysis (see below). In both cases, we ran simulations for 5000 units of time and then scored the outcome as polarized when the ratio of the maximum and minimum densities, assessed across all membrane compartments, was greater than the initial value.

#### Saturating positive feedback and no direct binding

We fixed values for the saturation factor 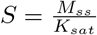 as specified in Fig. 5), and sampled values for all other parameters as in the case for linear positive feedback described above.

#### Model with mutual antagonism

For the model with mutual antagonism, we sampled values for *α* and *J_feedback_* uniformly between 0 and 1. We constrained the uniform steady state densities for proteins *A* and *B* to be *A_ss_* = 1, *B_ss_* = 1. We fixed a value for *J_A_* as indicated in figure panels. We sampled values for 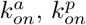, and 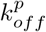 randomly between 0.1 and 1, then computed the values of all other parameters. We used LPA as described above to test for spontaneous and inducible polarization for the scenario in which diffusion is negligible.

### Relaxing the assumption of fast oligomerization kinetics

We used simulations of the full kinetic model to determine the effect of relaxing the assumption of fast oligomerization kinetics.

#### Linear positive feedback and no direct binding

We fixed 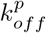 as indicated in (S2 Fig(A)), sampled values for all other parameters as described above for the one-species model, and tested for spontaneous symmetry breaking as described above.

#### Scenarios with direct binding

For the cases where *γ* = 1 and indirect positive feedback is active (*k_f_* ≥ 0), we fixed 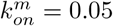 and 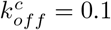; we assigned values to 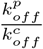 and *B_dir_* as indicated in (S3 Fig). We randomly sampled values for *J_feedback_* between 0 and 3 and *α* between 0 and 0.95. Then we computed the values of all other parameters. For the cases where *γ* < 1 and *k_f_* = 0, we fixed 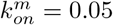 and 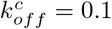; we assigned values to 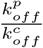 and *γB_dir_* as indicated in (S3 Fig). We sampled values for *J_feedback_* between 0 and 3 and *α* between 0 and 0.95. Then we computed the values of all other parameters. For both cases, we used numerical simulations to test for spontaneous polarization as described above.

### Relaxing the assumption that oligomers of size ≥ 2 do not dissociate from the membrane

To relax this assumption, we modified the full kinetic model so that oligomer dissociation rate decreases exponentially with size:

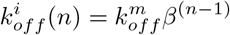

and the parameter *β* sets the rate of decrease. We fixed values for *β* as indicated in (S2 Fig(B)). We set 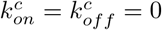, and 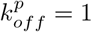. We sampled values for 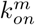 between 0.2 and 2, *k_f_* between 0 and 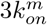 and 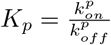 between 0 and ∞. Then we computed values for *J_feedback_* and mean oligomer size s at the uniform steady state, assessed by allowing simulations to reach steady state in a single spatial compartment. We then assessed spontaneous symmetry breaking as described above.

### Relaxing the assumption that oligomerization occurs only at the cell membrane

To relax this assumption, we modified the full kinetic model so that oligomerization occurs both in the cytoplasm and on the membrane with different equilibrium constants 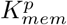 and 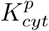. We assumed that oligomerization in the cytoplasm is at quasi-steady state, such that the distribution of cytoplasmic oligomer sizes is given by *C_n_* = *C*_1_*α*^*n*–1^ where 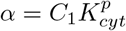. We set 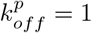 and we fixed the ratio 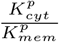 as indicated in (S2 Fig(C)). We sampled values for 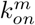 between 0.2 and 2, *k_f_* between 0 and 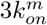, and *K_p_* between 0 and ∞. Then we computed values for *J_feedback_* and mean oligomer size s at the uniform steady state and tested for spontaneous polarization as described above.

### Using numerical simulations to determine polarization and boundary speed

#### Sampling parameter values

For the model with indirect positive feedback on monomer recruitment, we sampled values for *α* between 0.35 and 0.9 and *J_feedback_* between 2 and 10. We constrained *M_ss_* = 1, set the scaled monomer diffusivity to *D* = 0.00001 and sampled values for *K* between 0.2 and 2. To simulate the absence of size-dependent dissociation, we scaled 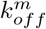 to hold the effective dissociation rate of molecules constant as *α* was varied. For simulations in which oligomerization dynamics were not assumed to be infinitely fast (S5 Fig), we assigned values to 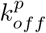 as indicated in the figure legend and assigned *J_feedback_* = 5. All other parameter values were determined by these constraints.

For the model with mutual antagonism, we sampled values for *α* between 0.35 and 0.9. We set *J_feedback_* = 0.64, 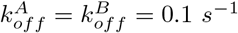, *J_A_* = *J_B_*, and 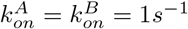, *D_A_* = *D_B_* = 0.1*μm*^2^ *s*^−1^, concentration of *A* and *B* were scaled to their uniform steady states respectively, and all other parameters were determined by these constraints. Simulations were initiated with one compartment 3-fold more concentrated than the uniform steady state. Polarization speed was assessed as the slope of the log of the maximum concentration at the membrane when the maximum concentration was 1.5-fold higher than its minimum value in time and 2-fold lower than its maximum value in time.

### Numerical simulations to determine boundary speed

In simulations to determine the boundary speed for the model with positive feedback, we sampled *α* between 0.35 and 0.8 and *J* between 2 and 10. Units for concentration, time, and length were scaled by *M_ss_*, 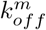, and membrane length respectively. Monomer diffusion was set to 0.00001 and 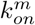 was randomly sampled between 0.2 and 2. If oligomerization dynamics were not assumed to be infinitely fast, 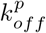 was set as indicated in figure panels. All other parameters were determined by these constraints. In simulations to determine boundary speed for the model with mutual antagonism, *α* was sampled between 0.35 and 0.8, *J_feedback_* = 0.64, 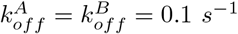, *J_A_* = *J_B_*, and 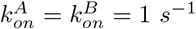, *D_A_* = *D_B_* = 0.1 *μm*^2^*s*^−1^, concentration of *A* and *B* were scaled to their uniform steady states respectively, and all other parameters were determined by these constraints. In simulations that lacked size-dependent dissociation, 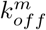 was scaled so that the effective dissociation rate of molecules off the membrane was held constant as *α* was varied. Simulations were initiated with an asymmetric step change distribution that was determined to be stable in the absence diffusion and where the polarized domain takes up 80 percent of the membrane length. For the model with positive feedback, this was determine by phase plane analysis (see below) and for the model with mutual antagonism this was determined through numerical simulations with *D_A_* = *D_B_* = 0. Boundary speed was determined by fitting a line to the position of the boundary over 400 time units following initiation of the simulation, defining the boundary as the inflection point of the spatial distribution.

### Phase plane analysis

To identify asymmetries involving polarized domains of defined sizes that are stable without diffusion, we performed phase plane analysis. We considered a model in which the cell membrane is divided into two membrane compartments exchange with a shared cytoplasmic pool, but we assumed no diffusive flux between membrane compartments.

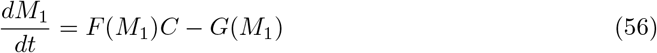

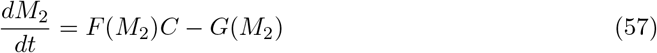

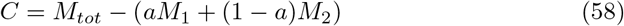

where *M*_1_ and *M*_2_ are the concentrations of protein in the two membrane compartments and *a* is the length membrane compartment *M*_1_ divided by the total length of the cell membrane. We plotted nullclines for these differential equations for defined values of *a* and used the polyxpoply function in MATLAB to determine intersections between the nullclines and thus steady states.

## Supporting information

**S1 Fig.**
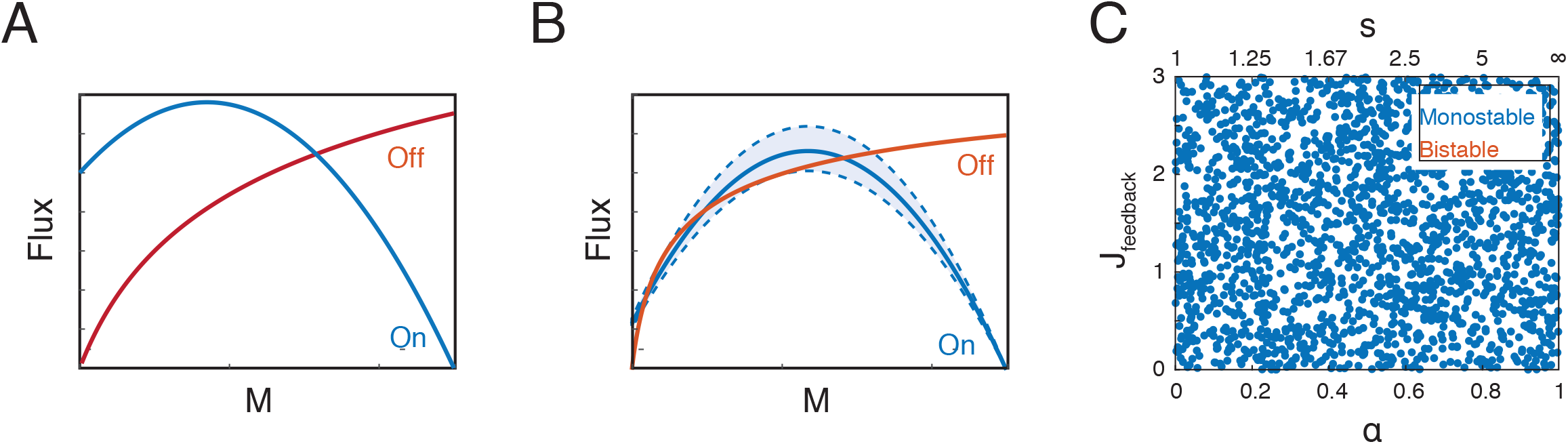
Steady state analysis of spatially uniform distributions. **(A)** Flux balance plot showing an example of a case in which there is exactly one uniform steady state. The blue line indicates flux of protein onto the membrane as a function of membrane protein concentration and the orange line indicates flux of protein off the membrane. **(B)** Flux balance plot indicating a narrow range of parameter values for which multiple uniform steady states are possible. Dotted lines indicate a range of curves that produce two uniform stable steady states, generated by varying *K*. **(C)** Scatter plot showing the existence of exactly one uniform steady state for the range of oligomerization strengths (*α*, *s*) and feedback strengths (*J_feedback_*) explored in this paper. Each point corresponds to a randomly chosen parameter set used in another analysis. Blue points have exactly one uniform steady state. There are no points that have multiple steady states.

**S2 Fig.**
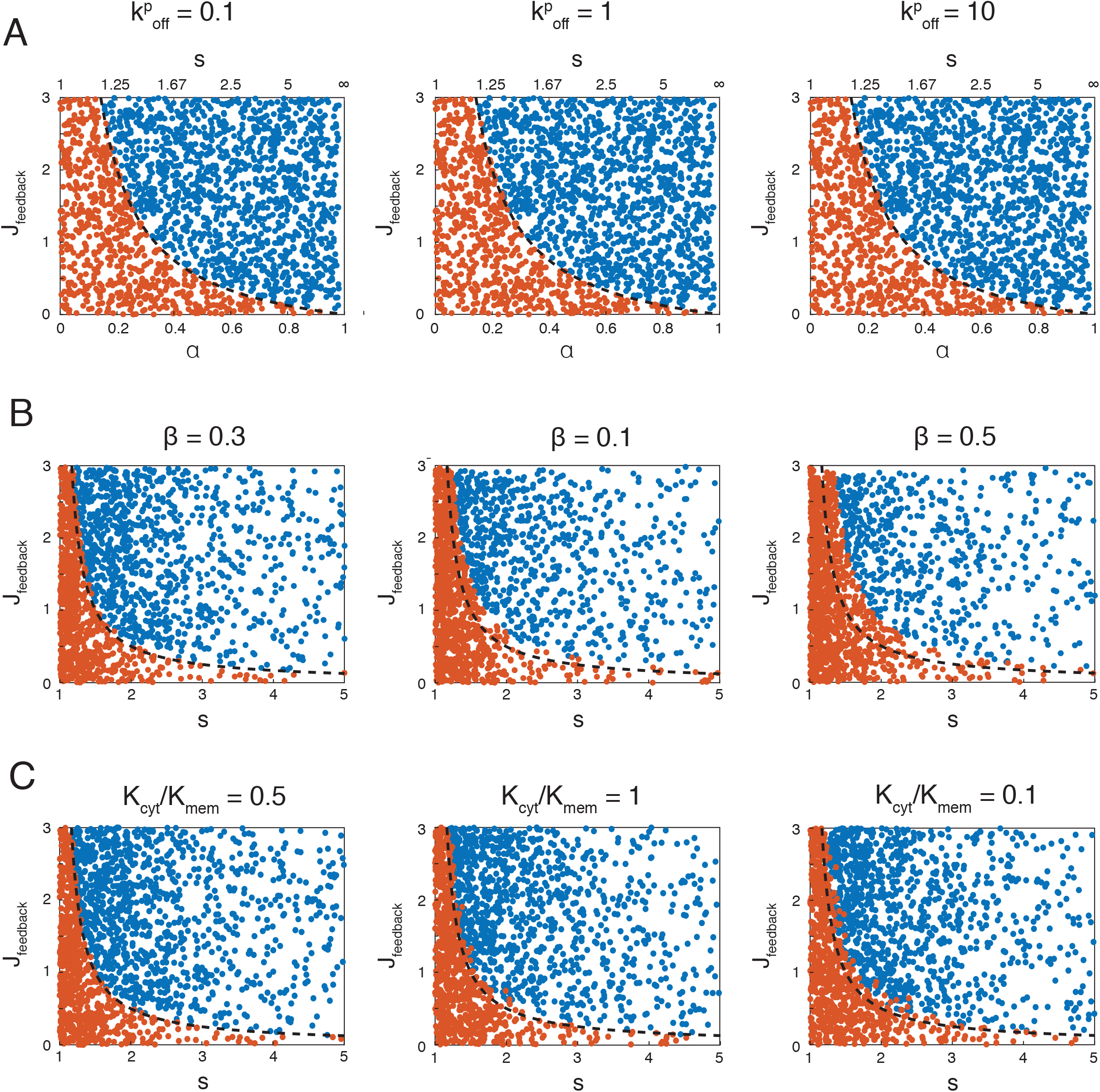
Effects of relaxing different assumptions on polarization. **(A)** Scatter plots showing results of simulations where the assumption that polymerization kinetics are fast relative to membrane exchange is relaxed. **(B)** Scatter plots showing results of simulations where the assumption that oligomers of size 2 and greater do not dissociated from the membrane is relaxed. In these simulations, the off rate of an oligomer of size *n* is equal to 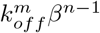. **(C)** Scatter plots showing the results of simulations where the assumption that oligomerization only occurs on the membrane is relaxed. *K_cyt_* denotes the equilibrium binding constant for oligomerization in the cytoplasm and *K_mem_* denotes the equilibrium binding constant for oligomerization at the membrane. In all panels, blue dots indicate spontaneous symmetry breaking, orange dots indicate a stable uniform state, and the dotted line indicates the predicted phase boundary between these outcomes based on the ideal case. Parameters were sampled as described in the methods section.

**S3 Fig.**
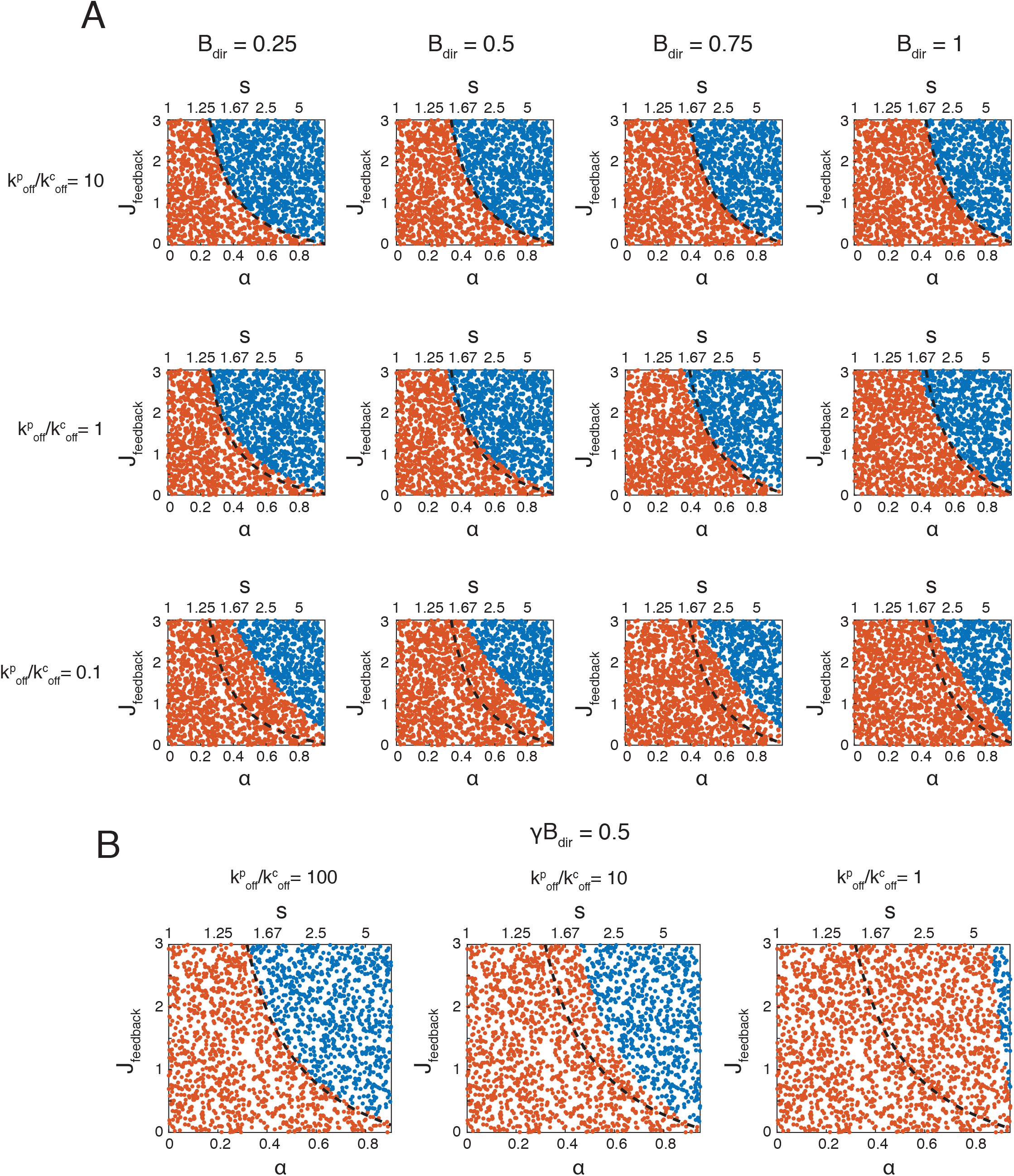
Predictions of the full kinetic equations for scenarios involving direct binding. **(A,B)** Scatter plots showing stability of the uniform steady state as a function of oligomerization (*α*, *s*) and feedback strength (*J_feedback_*) when **(A)**: *γ* = 1 for different values of *B_dir_* (left to right) and 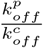 (top to bottom) or **(B)**: *γ* < 1 and *k_f_* =0 and *γB_dir_* = 0.5, for different values of 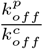 (left to right). Blue dots indicate unstable and orange dots indicate stable states. Dashed lines indicate the boundary between stable and unstable regimes derived for the one-species model.

**S4 Fig.**
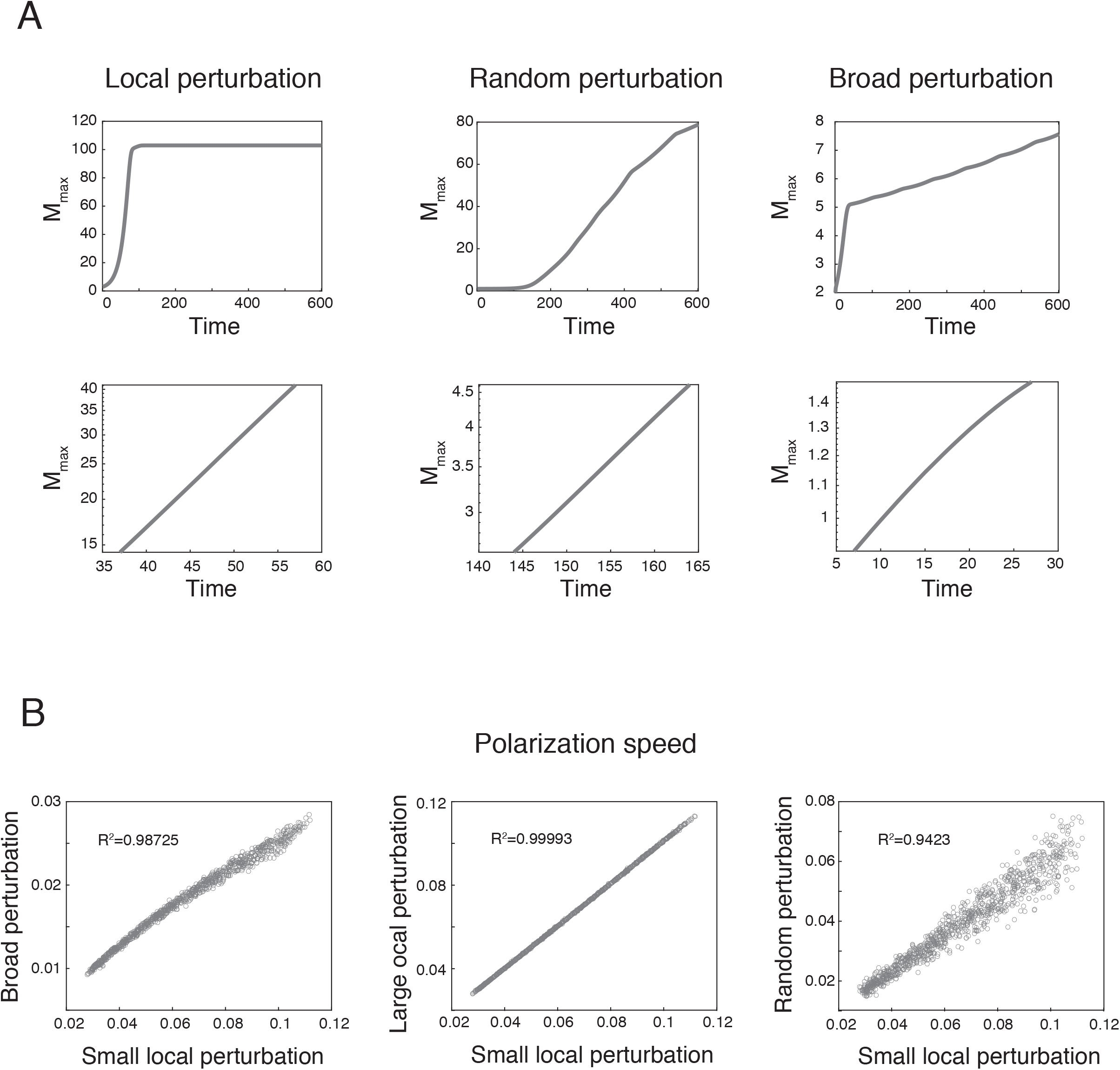
Exponential growth of a local peak is a reliable measure of polarization speed. **(A)** Plots showing examples of the logarithm of the maximum local value of *M* (*M_max_*) over time in simulations where polarization is triggered by different types of cues: a local perturbation, a spatially varying random perturbation, and a broad perturbation. While the speed of polarization differs based on the cue, all curves have a region in which *M_max_* grows exponentially. **(B)** Scatter plots showing the correlation between polarization speed across different types of perturbations in simulations where *J_feedback_* and *α* were sampled randomly. Each point corresponds to a unique choice of model parameter values.

**S5 Fig.**
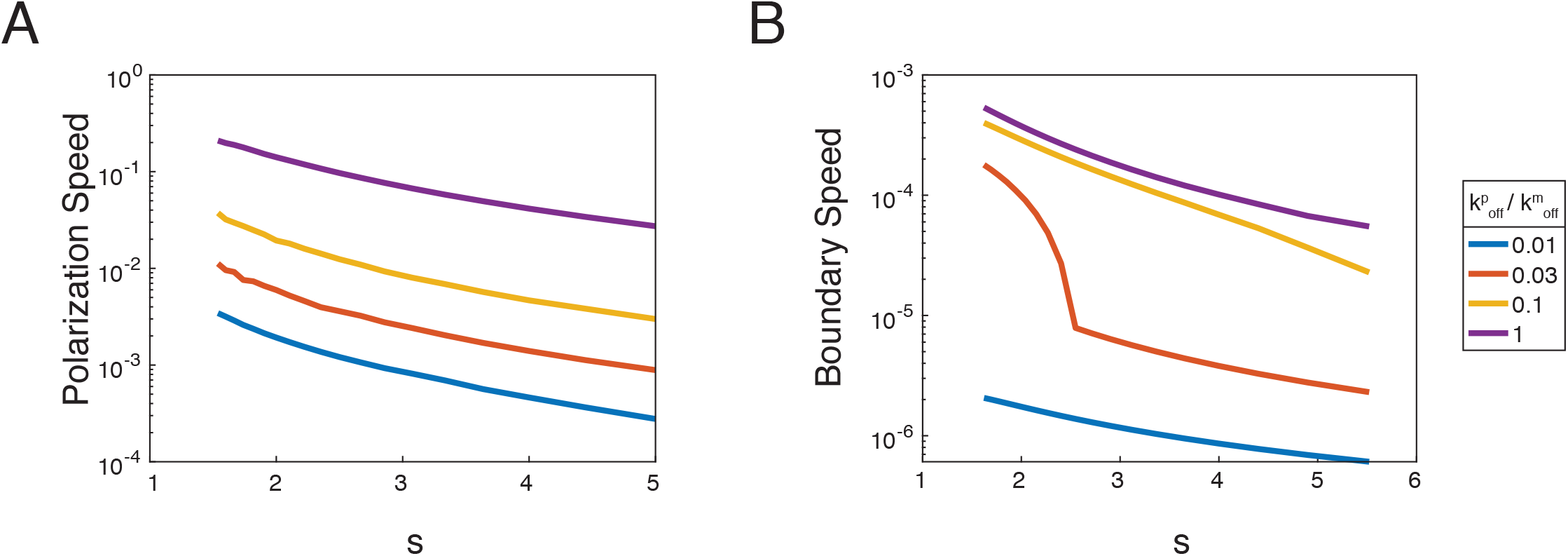
Dependence of polarization dynamics on oligomer dissociation kinetics. **(A,B)** Plots showing the effects of slowing down oligomerization dissociation kinetics 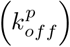 on polarization speed **(A)** and boundary speed **(B)**. *J_feedback_* is set to 5 in these simulations, while all other parameters are sampled as described in the methods. Colored lines show the dependence of polarization speed and boundary speed on mean oligomer size *s* for different values of 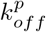.

**S6 Fig.**
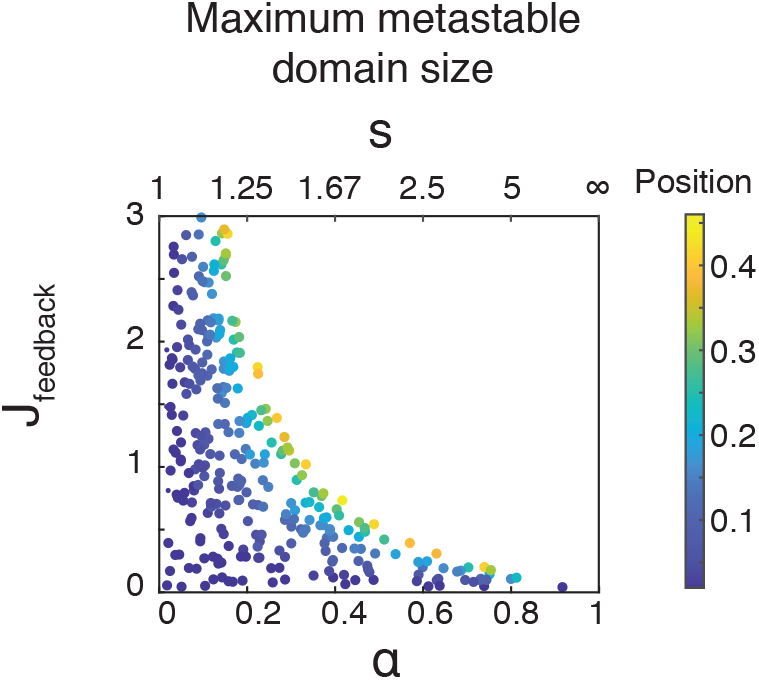
Asymmetries that do not arise spontaneously have a maximum metastable domain size. The maximum metastable domain size plotted as a function of *α* and *J_feedback_*. The color of individual points indicates the maximum size. Maximum metastable domain size was assessed using nullcline analysis assuming no diffusion and domain boundary position ranging from 50% to 2% cell length, evaluating at increments of 1% (see methods).

**S7 Fig.**
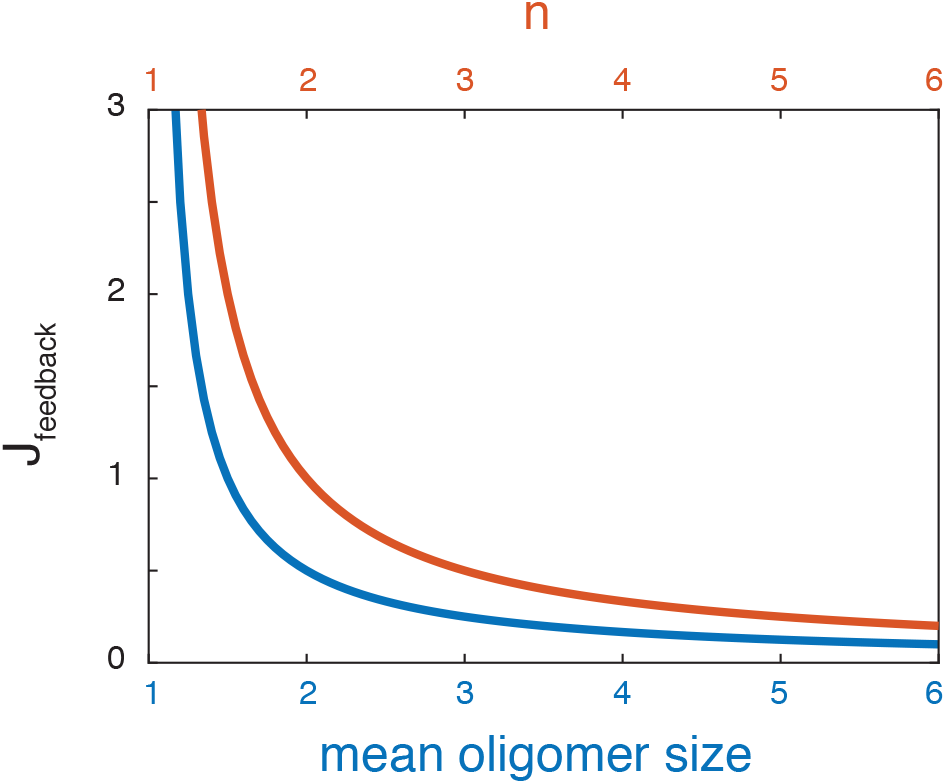
Feedback strength and the sharpness of non-linear dynamics determine spontaneous polarization for models with cooperative positive feedback. The orange line shows the boundary between spontaneously polarizing and stably uniform states for a model with higher order positive feedback in phase space defined by *J_feedback_* and the order of the feedback (n). The blue line shows the equivalent phase boundary for a model with linear positive feedback in phase space defined by mean oligomer size and *J_feedback_*.

## Acknowledgments

Work in E.M.M.’s lab was supported by the National Institutes of Health. Additionally, C.F.L. was supported by the National Institutes of Health (T32 GM007197).

## Notes

### Competing Interest Statement

The authors have declared no competing interest.

